# Spatiotemporal transcriptomic analyses reveal molecular gradient patterning during development and the tonotopic organization along the cochlear axis

**DOI:** 10.1101/2024.10.30.621022

**Authors:** Mengzhen Yan, Penghui Zhang, Yafan Wang, Haojie Wang, Junhong Li, Xiang Guo, Xiangyao Zeng, Zhili Feng, Zhiyi Yun, Fei Deng, Shouhong Wang, Di Deng, Lu Ma, Yong Feng, Huajun Wu, Yu Zhao, Jun Li

**Author notes:** Corresponding authors: Jun Li, Ph.D., Yu Zhao, Ph.D., Huajun Wu, Ph.D. Authors contributed equally to this work (Mengzhen Yan, Penghui Zhang, Yafan Wang, Haojie Wang).

## Abstract

While cochlear tonotopic organization is essential for sound frequency discrimination, the spatiotemporal molecular programs shaping this architecture remain unresolved. Here, we integrate spatiotemporal transcriptomics (E17.5-P8-adult) with single-cell RNA-seq to construct a single-cell-resolution atlas of mouse cochlear development. As marked by differential gradient expression genes, distinct spatially and molecularly defined hair cell (HC) subtypes along the apical-basal axis were identified: such as the apical Myo7a^+++^/Calb1^+++^ outer hair cell (OHC) subtype, the basal Myo7a^+^/Calb1^□^ OHC subtype, the apical vGlut3^+++^/Calb2^+^ inner hair cell (IHC) subtype, and the basal vGlut3^+^/Calb2^+++^ IHC subtype. In addition, the spatial data also demonstrate the developmental reversal of *Myo7a*/*Calb2Nefh* gradient patterns in HCs and spiral ganglion neurons (SGNs), revealing dynamic plasticity and complexity in frequency coding during development. Spatial analyses further demonstrate regional heterogeneity in cell communication intensity between HCs and SGNs, and also reveal core signaling hubs coordinating tonotopic organization, such as *Fgf10*-*Fgfr2* pathway in the basal region and *Ptn*-*Sdc* pathway in the apical region HC or SGNs at P8 stage. Our study provides an open-access spatial database and reveals the morphological and molecular foundations underlying cochlear tonotopic organization, linking molecular and developmental mechanisms to auditory pathophysiology.

## Introduction

The inner ear is one of the most structurally and functionally complex organs in vertebrates, comprising the cochlea and vestibular apparatus [1]. The cochlea is responsible for converting sound waves into neural signals, while the vestibular system aids in maintaining equilibrium [2, 3]. Development of inner ear is a highly coordinated process involving gene expression regulation, cellular differentiation, morphogenesis, and the activation of signaling pathways [3, 4]. The intricate architecture of inner ear is crucial for its sensory functions, and most genetic variations affecting its structure will cause sensory impairments. Cochlear and vestibular dysfunctions cause auditory and balance disorders, including congenital hearing loss, age-related hearing degeneration, and vestibular impairments, highlighting the need to elucidate inner ear development and function [3, 5]. Sensorineural hearing loss (SNHL) is one of the most prevalent chronic diseases, which typically affects high-frequency regions, though low and mid-frequency regions may also be involved depending on the etiology [6–8]. The cochlea exhibits a tonotopic gradient to sound frequencies along its longitudinal axis, from the base to the apex [9]. Hair cell (HC) loss or spiral ganglion neuron (SGN) damages are more predominant in the basal region under stresses, ototoxic drug exposure, or during aging [10–12]. However, the molecular basis underlying tonotopic organization and frequency-specific damage remain unclear, in part due to the complexity and inaccessibility of this deeply embedded organ [10, 13–15].

Single-cell transcriptomics have revolutionized our understanding of inner ear biology, including developmental processes [13, 16–18], age-related hearing loss (ARHL) [19], noise-induced hearing loss (NIHL) [20], and hair cell death/regeneration [21, 22]. This technology provides high-resolution insights into cell population heterogeneity within the inner ear, enabling the identification of novel cell types and constructions of developmental trajectories that were usually neglected in traditional bulk RNA sequencing approaches. For example, single-cell transcriptomics has identified rare cell types and transitional cell states in hair cell regeneration [22, 23] and novel SGNs subtypes in the cochlea [24–26]. These insights are invaluable for elucidating cellular contributions to inner ear structure and function, and for identifying potential therapeutic targets. However, single-cell transcriptomics has certain limitations [27], notably the loss of spatial context, which is essential for understanding inner ear function. For instance, single-cell transcriptomic studies have uncovered different SGNs subtypes [24, 26], however, their spatial distribution along the tonotopic axis remains unclear. Recent immunostaining studies have shown that type I SGNs exhibit differential distribution across cochlear regions [10], but the molecular mechanisms governing this phenomenon remain elusive, partly due to the loss of spatial information in single-cell RNA sequencing (scRNA-seq) data.

Spatial transcriptomics has emerged as a transformative approach, maintaining the spatial organization of cells while providing transcriptomic data [27]. Recent advances have enabled researchers to explore the spatial and temporal dynamics of gene expression in complex tissues like the heart [28, 29] and brain [30]. In the context of the inner ear, spatial transcriptomics allows for precise localization of gene expression, facilitating the study of tonotopic organization, development, and pathology. It also enables the integration of transcriptomic data with morphological findings, helping validate cellular identities in relation to specific tissue structures [29, 31]. Additionally, spatial transcriptomics offers insights into microenvironments and cell-cell interactions that drive organ development and function [32].

In this study, we aimed to establish the single-cell resolution spatial transcriptomic profile and decipher the molecular basis of the intricate architecture of cochlea, especially its tonotopic organization. By integrating spatial, single-cell transcriptomics, and RNA *in situ* validations, we found that a number of hair cell marker genes like *Myo7a* and neuronal marker genes like *Nefh* appeared in a gradient expression pattern along the cochlear axis. The gradient expression patterns of hair cell markers such as Myo7a, vGlut3 and Calb2 revealed the existences of HC subtypes and their finely tuned spatial arrangements. Furthermore, spatial differential gene expression analyses revealed regional heterogeneity of cell communication intensity between HCs and SGNs, with the weakest in the apex, aligning this region for detection of low-frequency sound. The differential signaling hubs were identified across different regions, such Ptn-sdc in the apical HCs and Fgf-Fgfr signaling pathway in the basal SGNs. These gradient patterns of critical marker genes, the regionalized heterogeneity of HCs or SGN subtypes, and spatially divergent cell communications are mirrored with the tonotopic organization along the cochlear axis. Notably, many gradient genes (e.g., *Myo7a*, *Fgf10*) are associated with deafness, implying their functional disruption directly impacts hearing. These findings offer valuable insights into the molecular mechanisms underlying the tonotopic organization of auditory sensitivity along the cochlear axis.

## Results

### Spatiotemporal characterization of cochlear cell types at different stages

To elucidate the molecular mechanisms of cochlear morphogenesis and its functional establishment during development, we conducted spatial transcriptomic analyses of the cochlea across three different important stages, including embryonic stage 17.5 (E17.5), postnatal stage day 8 (P8), and adult stage (2 months, 2M), respectively (Fig 1A). During development, the structures of the inner ear undergo dynamic changes, ultimately forming a complex and highly specialized architecture in adult stage (Fig 1A). Notably, the cochlea is a spiral structure rich in chambers, formed by various cell types that are intricately and spatially organized in an ordered manner (Fig. 1A) [1]. The spatiotemporal single-cell atlases were first established from single E17.5, P8 and 2M mouse cochlea section samples, respectively (Fig 1 and S1-S3 Fig).

**Fig 1.**
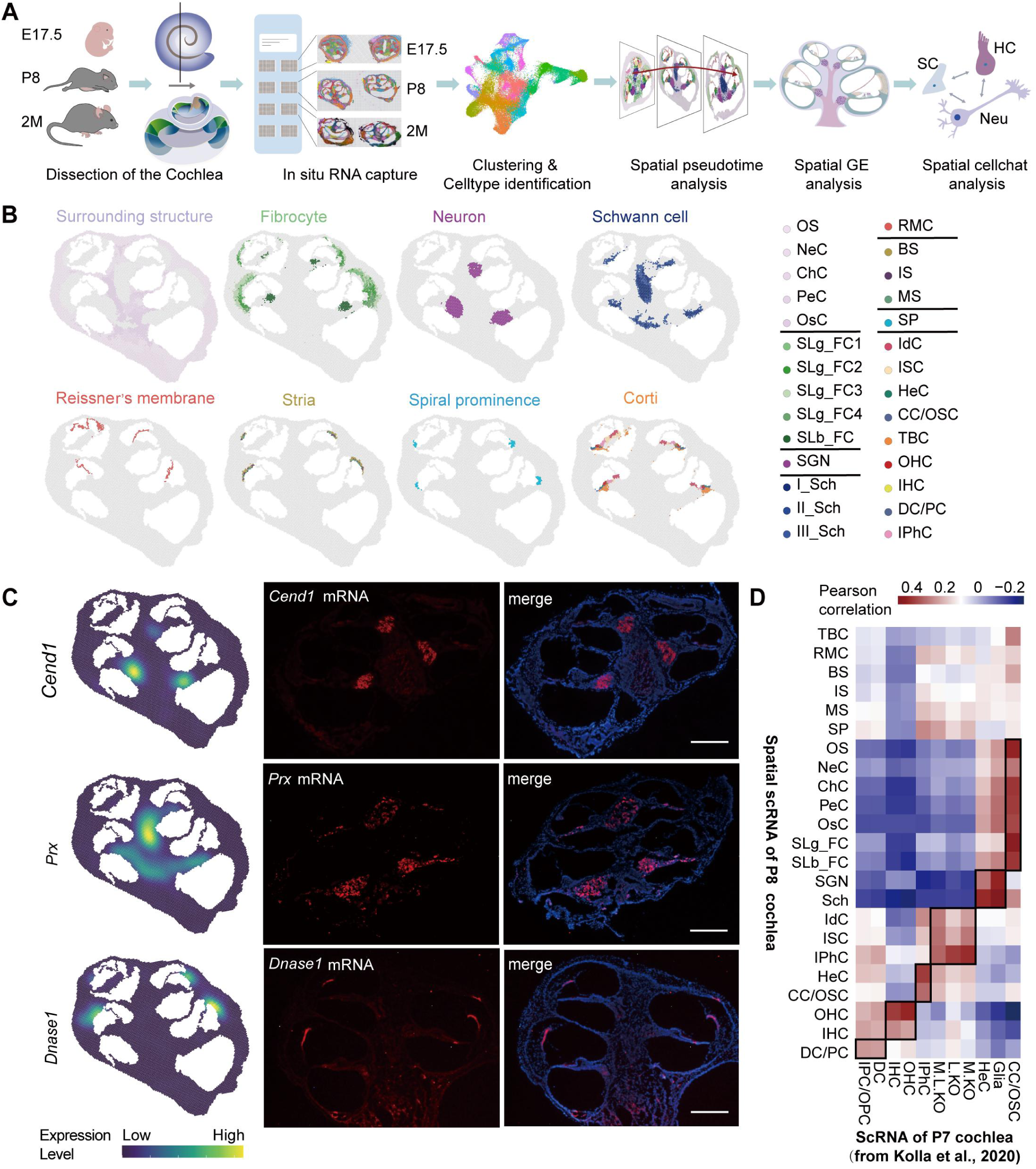
Spatial single-cell atlas of developing and adult cochlea. **(A)** The workflow of spatial transcriptomic assay for mouse cochlea at three stages (E17.5, P8, and 2M). **(B)** Spatial cross-section showing cell types localized in the P8 cochlea. Eight major groups of cell types were shown, respectively. **(C)** Spatial expression and RNA *in situ* detection of *Cend1*, *Prx* and *Dnase1* on the P8 cochlea cross-section. **(D)** Correlation heatmap showing the correspondence of cell annotation between spatial cells from P8 cochlea and the reported scRNA-seq dataset from P7 cochlea. OS: Out structure, NeuC: Neutrophils, ChC: Chondrocyte, PeC: Pericytes, OsC: Osteoblasts, SLg_FC1: Spiral ligament fibrocyte 1, SLg_FC2: Spiral ligament fibrocyte 2, SLg_FC3: Spiral ligament fibrocyte 3, SLg_FC4: Spiral ligament fibrocyte 4, SLb_FC: Spiral limbus fibrocyte, SGN: Spiral ganglion neuron, I_Sch: Type I schwann cell, II_Sch: Type II schwann cell, III_Sch: Type III schwann cell, RMC: Reissner’s membrane cell, BS: Basal stria, IS: Intermediate stria, MS: Marginal stria, SP: Spial prominence, IdC: Interdental cell, ISC: Inner sulcus cell, HeC: Hensen cell, CC/OSC: Claudius cell/Outer sulcus cell, TBC: Tympanic border cell, OHC: Outer hair cell, IHC: Inner hair cell, DC/PC: Deiter cell/Pillar cell, IPhC/IBC: Inner phalangeal cell/Inner border cell, M.KO:, Medial kölliker’s organ cell, L.KO: Lateral kölliker’s organ cell, M.L.KO: Medial-lateral kölliker’s organ cell, IPC/OPC: Inner pillar cell/Outer pillar cell.

In general, the spatial transcriptomic data have lower resolution and relatively low quality of cell clustering, compared to routine scRNA-seq data [27]. Integration of scRNA-seq and spatial RNA-seq will facilitate the spatial data analyses and enhance the powers of each other. Therefore, the E17.5 and P8 cochlear scRNA-seq data from previous publications [17], and the single-nucleus RNA-seq (snRNA-seq) data of the adult cochlear from our previous publication [33] were re-analyzed and compared with the spatial transcriptomic data. The gene markers from previous scRNA-seq studies [17, 24, 26, 33] were used for spatial clustering and cell type identification (S1 File). Totally, 23 cell types were annotated in E17.5 cochlea (S1-2A Fig), 28 cell types in P8 cochlea (Fig 1B), and 21 cell types in 2M cochlea (S1-3A Fig), respectively. Correlation analysis also revealed a strong consistency between our spatial scRNA datasets and previously published related scRNA datasets (Fig 1D, S1-2C and S1-3C Fig).

According to the spatial distribution, all cell types from each dataset were briefly split into 8 groups including surrounding structure, fibrocyte, neuron, Schwann cell, Reissner’s membrane, stria, spiral prominence, and organ of Corti (OC) (Fig 1B, S1-2A and S1-3A Fig). Neither spiral prominence was identified in the dataset of E17.5 cochlea, but they were clearly identified in P8 and adult cochlea single-cell atlases with markers *Aqp5* and *Bmp6*, suggesting these cells were differentiated cell type in postnatal stage (S1-1A, S1-2B and S1-3B Fig). The surrounding structure of P8 cochlea mainly contained outer structure, identified with *Acp5* and *Mmp9* gene markers, Neutrophils with *S100a8* and *S100a8* gene markers, chondrocyte and pericytes with *Penk*, *Igfbp6*, *Acta2* and *Tagln* gene markers, and Osteoblast with *Ifitm5* and *Sgms2* gene markers (S1-1A Fig). Fibrocytes of cochlea mainly distributed in the spiral ligament (SLg) and limbus (SLb) [33, 34], and were discovered in all three datasets, identified by *Car3* and *Otor* in P8 dataset (S1-1 Fig). The neuron cells were found mainly existed in the three regions near the cochlea duct with the known neuron markers (*Nefh*, *Snap25*, *Nefl* and *Meg3*) (S1-1, S1-2B and D, S1-3B and D Fig and S1 File) [35–37]. Spatial expression and RNA *in situ* detections supported *Cend1* could also be one of the markers to identify SGNs cluster. Spatial analyses also identified Schwann cells and their subtypes, which serves as the supporting cells of SGNs (Fig 1B and S1 File), identified by *Prx*, and *mpz* (Fig 1C, S1-1 Fig), suggesting their role in myelination processes [38]. Distinct cell types identified in the OC revealed the high cellular heterogeneity during hearing establishment and maturation, such as 7, 9 and 6 cell types in E17.5, P8 and adult samples (Fig 1B, S1-2A and S1-3A Fig), respectively. In the OC of P8, interdental cell with gene marker *Otoa* and *Rspo2*, inner sulcus cell with markers *Clic6*, inner phalangeal cell with markers *Tectb* and *Tecta*, Hensen’s cell and outer sulcus cell with markers *Frzb*, *Npnt* and *Bmp4*, outer hair cell (OHC) and inner hair cell (IHC) with markers *Otof*, and Deiter cell/pillar cell with markers *Fgfr3* were observed (S1-1 Fig).

After brief identification of cochlear cell types, several specific cochlear structures and cell types were further spatially analyzed in details to validate the spatial scRNA atlas. The lateral wall, an essential component of the scala media, includes the stria vascularis, which plays a crucial role in balancing the endocochlear potential, and the spiral ligament, which provides structural support and participates in ion homeostasis (Fig 1B) [34]. There were three cell types in the location of stria vascularis from E17.5 and P8 samples (Fig 1B and S1-1A Fig), including basal stria, intermediate stria and marginal stria that were identified by marker genes *Dnase1*, *Tyr* and *Kcne1*, respectively (Fig 1C, S1-1A and S1-2B Fig). *Dnase1* was specifically expressed in the stria vascularis (Fig 1C), which was consistent with their expression profiles revealed by spatial data analyses. But only basal and marginal cells were annotated in the adult dataset, using *Dct*, *Enpep* and *Kcne1* markers (S1-3B Fig). The intermediate cells were not identified, probably due to ossified structures in adult cochlea which hinders the tissue sections and detection of mRNA. Spiral prominence cells were identified between outer sulcus cell and stria vascularis with specific high expression of *Aqp5* (S1-1 Fig), which was consistent with previous cochlear snRNA-seq study [33]. In addition, four types of fibrocytes were identified in the lateral wall, based on spatial location and known markers (*Car3*, *Otor*) (Fig 1B, S1-1A-B Fig and S1 File) [20]. Above results revealed a strong correspondence between the spatial data and the single-cell level RNA-seq data, supporting the accuracy and high resolution of our spatial single-cell atlas. Furthermore, our spatial transcriptomic datasets enabled better identification of diverse cell types within complex cochlear structure and provided insights into the spatial transcriptional changes.

### Spatiotemporal dynamics and gradients of spiral ganglion neuron subtypes

Spatial analyses further identified SGNs the primary neurons transmitting signals from HCs (Fig 2A) in their exact anatomical regions. To explore the molecular changes that occur within the neuronal differentiation and maturation, we fitted a principal tree in diffusion space to show the temporal dynamics of developing SGNs (E17.5, P8, and adult) (Fig 2A and B). The tree was represented into a dendrogram recapitulating a branched trajectory based on the transcriptional similarity of pseudotime-ordered cells (Fig 2B). We further used differential gene expression (DEG) approaches that characterize pseudotime-ordered molecular trajectories (Fig 2C). Focusing on genes with significant variations (Fig 2C), we conducted gene ontology analysis to uncover different biology processes during SGNs development and maturation (Fig 2D). we observed that the shared, unspecialized neurons’ trajectory was characterized by rapid downregulation of genes *Ina*, *Fabp7* and *Tubb2b* commonly associated with early neuronal differentiation processes, and upregulated of genes *Calb2*, *Snap25* and *Spp1* that are involved in specification events in cochlea and synapse function (Fig 2D and E, S2A Fig). Also, branched expression analysis modeling (BEAM) was conducted to identify genes showing significant expression differences at cell fate branch points (S2B Fig), and GO analysis was further employed to illustrate the related function (S2C Fig). The bubble diagram, pseudotime gene expression plots and bifurcation gene expression plots more visually illustrated the dynamics of these genes during development (Fig 2E, S2A and D Fig). The differential marker gene expression analyses revealed the relatively higher expression of *Calb2*, *Snap25*, *Spp1* and *Nefm* in adult neurons, while genes such as *Fabp7*, *Tubb2b*, *Ina* and *Tubb3* showed relatively higher expression at earlier stages (Fig 3E, S2A Fig), highly related to neuron system development, axon formation and cytoskeleton organization (S2C Fig). This temporal dynamic gene expression pattern of SGN was further validated by immunostaining with Calb2 and Calb1 (Fig 2F). At E17.5 stage, the highest expression level of Calb2 was detected at the apical region, medium at the middle region, and weak at the basal region (Fig 2F). This pattern was gradually changed at P8 stage (Fig 3F), and finally reversed in the adulthood, with the highest expression level of Calb2 in the basal SGNs (Fig 2F). These spatiotemporal dynamics of gene expressions reflected the molecular mechanism of development and maturation of SGNs from E17.5 to adulthood.

**Fig 2.**
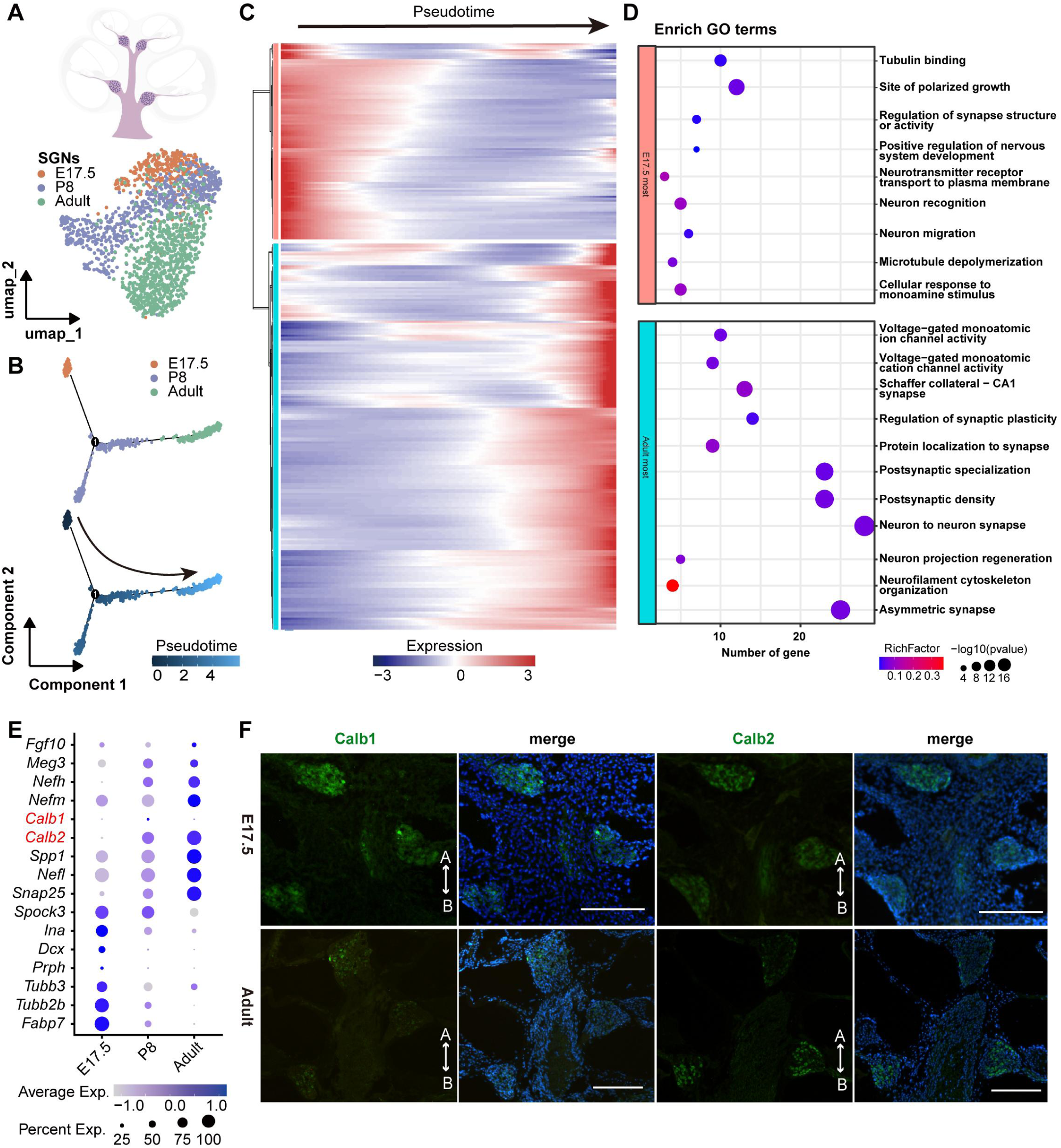
Spatiotemporal dynamics of gene expression in spiral ganglion neurons. **(A)** Upper, the diagram of neuronal system in cross-section of cochlea. Bottom, the umap spot showing the neuron cells extracted from E17.5, P8 and adult cochlea. **(B)** Plots recapitulating the branched trajectory of the developing SGNs (E17.5, P8, and adult) based on the transcriptional similarity of pseudotime-ordered cells. **(C)** Heatmap showing the pseudotime analysis for neuron cells from E17.5, P8 and adult dataset, with the top 10 upregulated genes along the differentiation of each branch. **(D)** Gene ontology analysis of differential genes from **(C)**. **(E)** Bubble plot showing the differential expression of a set of genes in neuron cells at different development stages. The size of the dots represents the percentage of cells in the clusters expressing the genes and the color intensity represents the average expression levels of the gene in that cluster. **(F)** Immunostaining of Calb1 and Calb2 (green) on the E17.5 and P8 cochlea cross-sections. DAPI is labeled in blue. A, apex; B, base. Scale bar equals to 200 μm.

**Fig 3.**
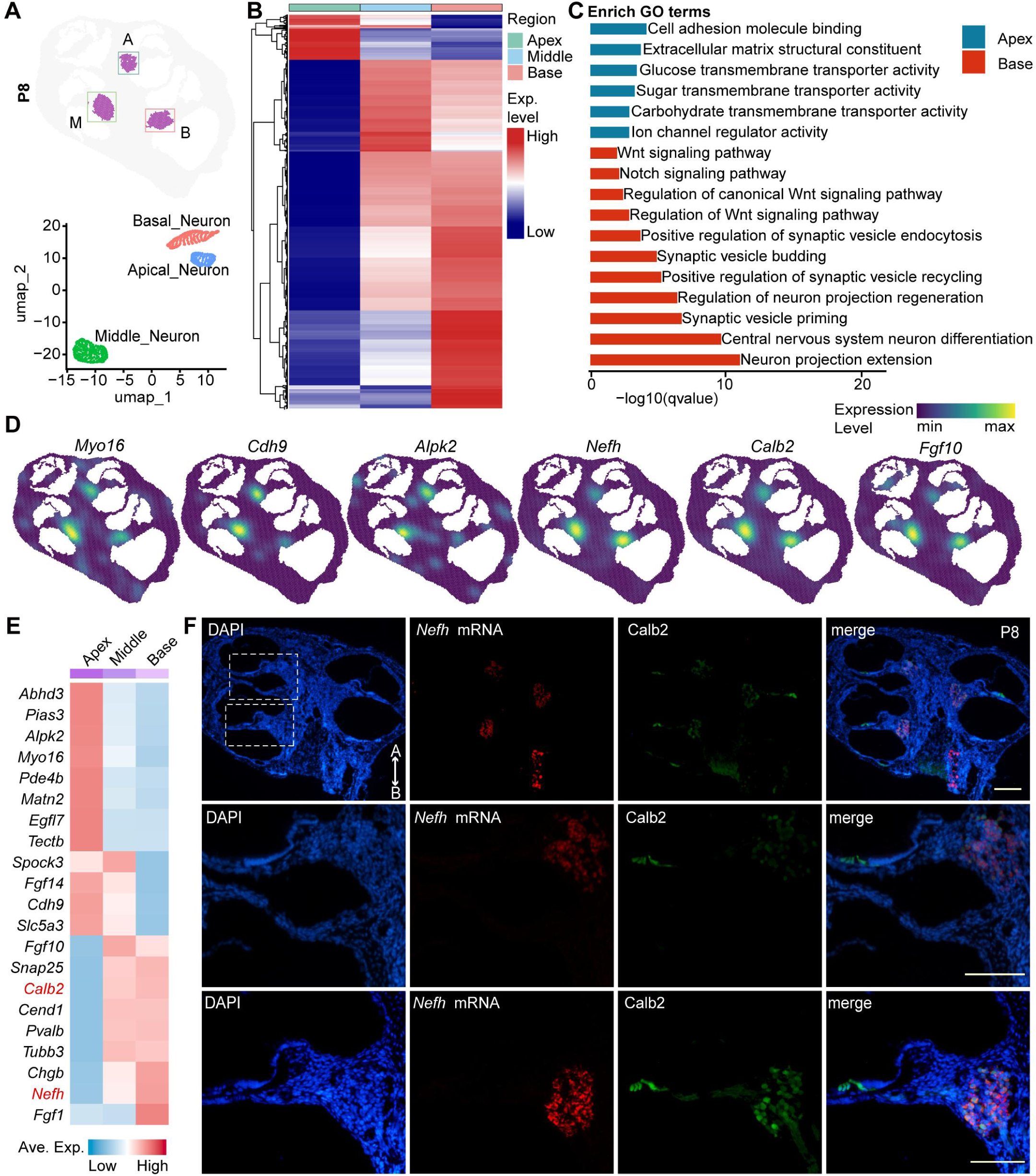
Spatial differential analysis reveals gradients of neuronal gene expression from apical to basal SGNs. **(A)** Upper, spatial painting of spiral ganglion neurons (SGNs) at P8 cochlea. The rectangular boxes with different colors indicate three different regions. A, apical; M, middle; B, basal. Bottom, UMAP plot showing the apical middle and basal neurons extracted from P8 cochlea. **(B)** Heatmap showing the expression level of the gradient genes representing two patterns in P8 SGNs (decreasing or increasing expression pattern from apical to basal regions). **(C)** Gene ontology analysis of differential genes from **(B). (D)** Density plots showing the spatial expression patterns of *Myo16*, *Cdh9*, *Alpk2*, *Nefh*, *Calb2*, and *Fgf10* on P8 cochlea cross section. **(E)** Heatmap showing two expression patterns of the 21 functional gradient genes in SGNs (decreasing or increasing expression pattern from apical to basal regions). The genes which were confirmed by RNA *in situ* or immunofluorescent staining assay were labeled in red. (**F**) Immunostaining of Calb2 (green) and RNA *in situ* detection of *Nefh* mRNA (red) on P8 cochlea cross-section. DAPI is labeled in blue. Magnified views of stained neuron cell in the apex region and basal regions were shown in the middle and bottom rows, respectively. A, apical; B, basal. All scale bar equals to 200 μm.

**Fig 4.**
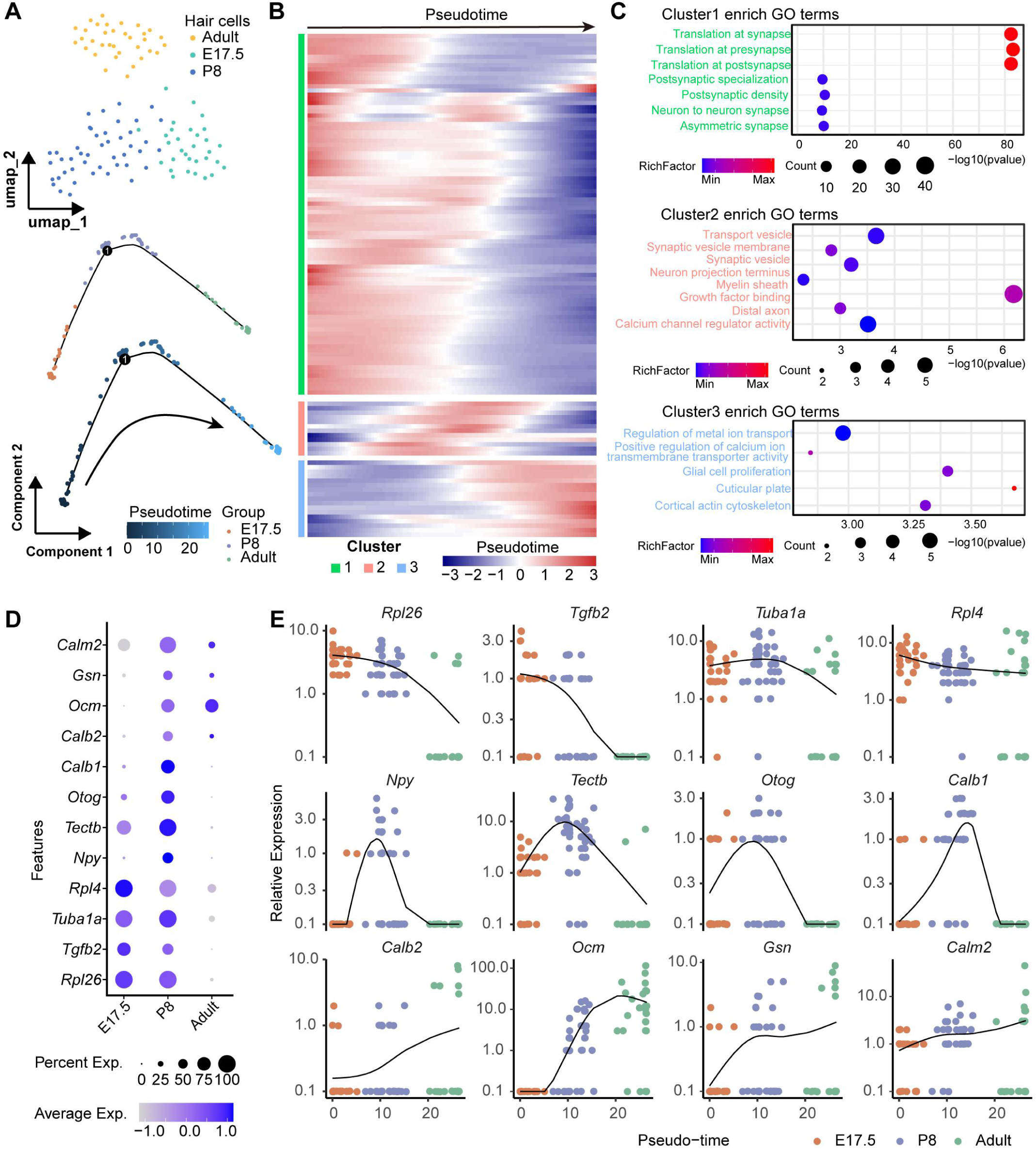
Spatiotemporal analysis for the hair cells. **(A)** Upper, the umap spot showing the hair cells (HC) extracted from E17.5, P8 and adult cochlea. Bottom, pseudotime trajectory plot showing the developing HC (E17.5, P8, and adult) based on the transcriptional similarity of pseudotime-ordered cells. **(B)** Heatmap showing the transcriptional changes for hair cells from E17.5, P8 and adult dataset along pseudotime. **(C)** Gene ontology analysis of differential genes from clusters in **(B)**. **(D)** Bubble plot showing the differential expression of 12 functional genes in HC at different development stages. The size of the dots represents the percentage of cells in the clusters expressing the genes and the color intensity represents the average expression levels of the gene in that cluster. **(E)** Curve plots showing the expression changes of the functional genes in developing HC along pseudotime.

To further comprehensively investigate the spatial differences of SGNs, we performed local single-cell analyses from apical to basal regions of cochlea. Neuron cells from three distinct regions (apical, middle, and basal) were isolated according to cell identifications and their spatial coordinates from the total spatial single-cell atlas (Fig 3A and S3-1A, B Fig, left).

After pairwise comparisons among apical, middle, and basal SGNs (Fig 3B and S3-1A, B Fig, right), the obtained differentially expression genes (DEGs) were intersected with the neuron-specific gene list from snRNA-seq dataset and spatial dataset (S1 File). Finally, 323, 470 and 1587 genes were identified from E17.5, P8, and adult datasets, respectively (S2 File).

For E17.5 spatial dataset, *Nefm*, *Nefl*, *Trim2*, and *Sema3a* genes were highly expressed at the apical SGNs and relatively lower in the basal SGNs (S3-1C and E Fig). *Fgf11* and *Abr* genes showed opposite expression pattern (S3-1 C and E Fig). Two obvious contrary expression patterns were maintained at P8 stage (Fig 3B, D and S2 File). Genes like *Nefh*, *Calb2*, and *Fgf10* were expressed higher at basal region, but *Myo16*, *Cdh9*, *Alpk2* and *Tectb* were expressed higher at the apical region (Fig 3D and E). GO analysis showed that the apex-high genes at E17.5 and P8 were related with protein binding, molecule transporter activity while the base-high genes mainly were involved in Neuron development and synapse formation (Fig 3C and S3-1F Fig). Expectedly, the apex-high genes at adult were associated with neuron function and the base-high genes were related to the function of Neuron, synapse and cykeleton organization. RNA *in situ* and immunostaining with these gradient markers *Nefh* and Calb2 were performed to validate the gradient expression pattern (Fig 3F). *Nefh* mRNAs are most abundantly detected at the basal region (Fig 3F, bottom), and weak at the apical region (Fig 3F, middle). Calb2 antibody immunostaining also displayed similar pattern, but the variation of gradient was not as sharp as *Nefh* (Fig 3F) at P8 stage. Note that both *Nefh* and *Calb2* expression patterns were same at E17.5 and P8 stage (S3-1D Fig and Fig 3D, E). However, *Nefh* and *Calb2* were expressed at a higher level in the apical SGNs and lower in the basal region at the adult stage (S3-1D Fig and G Fig). Furthermore, according to SGN markers from the published data [24, 26, 33], *Nefh* was commonly expressed in all SGN subtypes but with higher expression level in subtype III [33]. The relative higher expression of *Nefh* in basal region suggested that SGN subtype III may be predominantly located at the basal region. The expression of other gene markers suggested that SGN subtypes may exhibit a similar gradient distribution along the cochlear axis, based on their marker gene expressions (Fig 3B and D, and S3-1 Fig). These spatiotemporal dynamics of neuronal gene expression gradients and distributions of SGN subtypes revealed the progress of SGN development and differentiation along the cochlear axis (Fig 3C and S3-1F Fig).

For adult cochlea, Type I and type II are two main subtypes, constituting of 90-95% and 5%-10% in the SGNs, respectively. Type I SGNs were further divided into subtype Ia, Ib, and Ic [35, 37]. An additional subtype, type III was discovered in our recent study (S1-3B Fig) [33]. Due to close similarity of each SGN subtype, it is challenging to further identify each SGN subtype from these spatial sequencing data. Therefore, immunostaining and RNA *in situ* detections with several SGN markers were performed to identify each SGN subtype (Fig. 3C-E). A gradient labeling of Calb1^+^ type 1b SGNs was observed in whole amount adult cochlea or sections (S3-1E and S3-2A Fig). The Calb1 was relatively higher expressed in the apical SGNs. For Pou4f1^+^ type Ic SGNs, relatively more cells were found in the apical and middle regions, less in the basal region (S3-2D Fig). The Calb2 was dominantly expressed in type III SGNs [10] and its expression exhibited a basal-to-apical gradient in the whole amount and section cochlea (S3-1G and S3-2B Fig). The type II SGN marker genes *Cilp* exhibited relative stronger expression levels in the middle or basal adult SGNs in adult cochlea (S3-2C Fig). These results confirmed the diversity and spatial dynamics of SGN subtypes in adult cochlea.

### Spatiotemporal profiling of the organ of Corti development

The OC is embedded in the cochlear duct and rests on the basilar membrane (Fig 5A, left), allowing it to be finely tuned to different frequencies of sound [40]. Spatial visualization demonstrated distinct OC cell types at adult, including inner and outer hair cells (IHCs and OHCs), Deiter cells/Pillar cells, Hensen’s cells, tympanic border cells, and interdental cell (Fig 5A, right). The expression profiling of key marker genes across various identified cell clusters in the snRNA-seq dataset (S1-3B Fig, right), consistent with the spatial transcriptomics findings (Fig 5A and S1-3B Fig), reinforcing the identification of these cell types. For example, *Slc17a8* was predominantly expressed in IHCs, while *Ocm* was highly expressed in the OHCs (Fig 5B). Overall, the consistent patterns observed between spatial transcriptomics and scRNA-seq data confirmed the accuracy of cell type identification and their respective marker genes, providing a comprehensive view of cochlear cell diversity.

**Fig 5.**
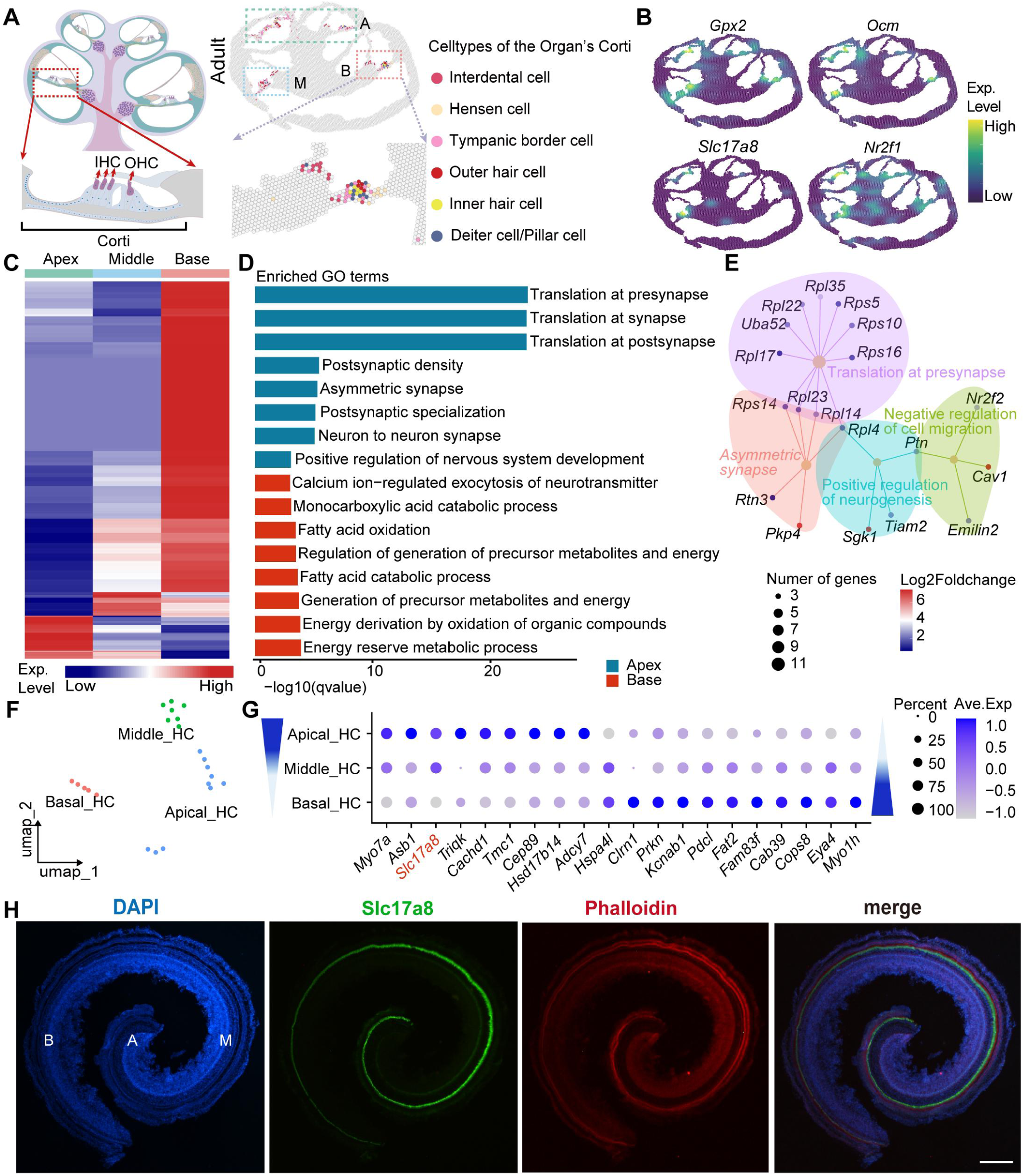
Spatial differential analysis reveals gene expression gradients of Corti organ from apex to base. **(A)** Left, the diagram of cochlea cross-section and magnified view of the cells from the organ of Corti (OC). Right, spatial painting of cell clusters from OC identified in adult cochlea and the magnified view to show the cells in OC. **(B)** Density plot showing the spatial expression of several cell-type marker genes of OC. **(C)** Heatmap showing the expression patterns of the gradient genes representing two patterns in P8 SGNs (decreasing or increasing expression pattern from apical to basal regions). **(D)** Gene ontology analysis of differential genes from **(C). (E)** Network plot showing several significant GO terms and its related genes enriched in the apical adult OC. **(F)** Umap spot showing the HCs extracted from different regions in the adult cochlea spatial section. **(G)** Bubble plot showing the gradient expression patterns of 20 functional genes in HCs at different regions. The size of the dots represents the percentage of cells in the clusters expressing the genes and the color intensity represents the average expression levels of the gene in that cluster. **(H)** Whole-mount immunostaining of Slc17a8 (green) and phalloidin (red) in adult cochlea. DAPI is labeled in blue. Scale bars equal to 200 μm.

To further understand the changes of expression pattern occurring in the process of hair cell maturing, we fitted a master tree for hair cells from the three stages in diffusion space. The tree was then represented as a dendrogram that reproduced a branching trajectory based on the transcriptional similarity of the proposed time-ordered cells (Fig 4A). The generated tree accurately reflected the developmental stage branching characteristics. Dynamic gene expression changes across the three stages were demonstrated by heatmaps (Fig 4B), with genes such as *Ocm* and *Tgfb2*, displaying distinct temporal expression patterns (Fig 4D). Genes such as *Rpl26*, *Tectb*, *Otog*, *Npy*, and *Calm2* exhibited dynamic expression from E17.5 to adulthood, confirming transcriptional changes during cochlear development (Fig 4D). The proposed temporal trajectories of key genes were demonstrated, illustrating the dynamics of cellular expression during developmental stages (Fig 4E). *Rpl26*, *Tgfb2*, *Ocm*, and *Gsn* exhibited stage-specific expression peaks during hair cell maturation (Fig 4E). The GO analysis further revealed the functional shift underlying hair cell development (Fig 4C). Overall, this analysis revealed the key spatiotemporal gene expression features within the HC that accompany the maturation of cochlear cell types and supporting structures critical for hearing function.

### Spatiotemporal gradient patterning of hair cell markers reveals the existence of IHC and OHC subtypes

The hair cells (HCs) were defined around E16.5 stage following a basal-to-apical gradient [41]. All the HCs were well differentiated at E17.5, including the apical regions with relative weaker expressed marker *Atoh1* and *Pou4f3* (S1-2B Fig, right). No significant differences of Myo7a signals were observed between IHCs and OHCs from apex-to-base. At P7 stage, the Myo7a expression in apical region was clear and became stronger compared to E17.5 cochlea, revealing further differentiation of hair cells in the apex. The expression of Myo7a in the middle region was relatively stronger than that in the apical and basal regions (S4B Fig). In adulthood, the Myo7a expressions in IHCs display an averaged pattern from apex-to-base, relatively slightly higher in the middle region (S4C and D Fig). While the Myo7a expression in OHCs showed a huge divergence from apex-to-base (S4C and D Fig). The basal OHCs expressed very low level of Myo7a (S4C and D Fig). It’s worth noting that the sharp contrast of Myo7a signals between IHCs and OHCs in the basal region (S4 D Fig). At the middle region, the expression of Myo7a was gradually increased in OHCs following a basal-to-apical gradient, basal Myo7a^+^ OHCs vs apical Myo7a^+++^ OHCs (S4 D Fig). The Myo7a signals of OHCs and IHCs showed less divergence along the gradient and became no obvious divergence at the apex (S4 D Fig).

To further validate this expression gradient in Corti, a number of spatial differentially expressed genes (DEGs) were isolated from E17.5 (S5-2A Fig, left), P8 (S5-2A Fig, right), and adulthood (Fig 5C). Simultaneously, GO analysis demonstrated that the apex-high genes (like *Cyp26a1*, *Nr2f2*, *Fst*, *Sox10*, and *Six4*) in E17.5 and P8 OC both played a critical role in Wnt, BMP, and RA signaling pathways, while the base-high genes at E17.5 mainly were enriched in development related GO terms like inner ear development and connective tissue development (S5-2B and C Fig). However, the base-high genes at P8 were involved in the function of ion channel activity and cell differentiation (S5-2C Fig). At adult, the genes (like *Ptn*, *Cav1*, *Rps14*, and Emilin2) apically enriched in OC largely involved in the synapse function specialization and the base-high genes were associated with the function of energy metabolic process (Fig 5D, E). The above results revealed the OC development involved coordinated processes of structural and functional maturation. After filtering and intersecting with a dataset of HC-specific genes (S1 File), 184, 265, and 205 DEGs were retained for the three datasets, respectively (S3 File). There are two spatial patterns of gene expression gradients for these spatial DEGs. In adulthood HCs, genes like *Myo7a*, and *Tmc1* displayed a decreasing gradient from apical to basal pattern, whereas genes *Myo1h*, *Eya4*, *Clrn1*, *Prkn*, and *Kcnab1* showed a reversed gradient in HCs (Fig 5 F and G). The relatively higher expression of damage protection-related genes *Hspa4l*, *Clrn1*, *Prkn*, and *Kcnab1* in the basal HCs in contrast to the apical HCs suggests a greater need for protection from damage, explaining the vulnerability of basal HCs in high-frequency hearing loss.

These spatial DEG data, particularly the *Myo7a* gene, were well matched with Myo7a immunostaining results in the whole amount (Fig 5G and S4C Fig) of adult cochlea. Apical OHCs exhibited comparable Myo7a expression levels to IHCs, while basal OHCs showed lower expression (S4D Fig). This gradient, developmentally dynamic at the E17.5 (S4A Fig) and P8 stages (S4B Fig), stabilized in adulthood. Similar or reversed gene expression gradients were also found in the E17.5 and P8 cochlear HCs, including *Calb1*, *Tmc1*, *Eya1*, and *Otof* genes (S5-3A-D Fig and S3 File). The spatial DEG analyses and the validation by immunostaining both provided compelling evidence that the adult HCs from apex to base are spatially distinct subtypes, as revealed by their different or even reversed gradient patterns of HC marker gene expressions. The Calb1 positive HCs are only found at the apical region (S5-1A Fig). In contrast, vGlut3 (Slc17a8) showed higher expression in the apical and middle IHCs (Fig 5H). The transcription factor Foxp1 was strongly expressed in the middle OHCs, less in the basal, and few or none in the apical OHCs (S5-1B and C Fig). Immunostaining with Prestin (Slc26a5, the specific known marker of the OHCs) antibody revealed that its expression exhibits a relatively averaged expression level in adult OHCs across the cochlear axis (S5-1D Fig), while an increasing expression from apex to base in P8 OHCs (S5-3E and F Fig). Based on these validated gradient markers, we defined the OHCs and IHCs as at least four distinct subtypes: the Myo7a^+^/Calb1^□^ basal OHC subtype, the Myo7a^+++^/Calb1^+++^ apical OHC subtype, the vGlut3^+^/Calb2^+++^ basal IHC subtype, and the vGlut3^+++^/Calb2^+^ apical IHC subtype, respectively. These gradient patterns of OHC and IHC subtypes with differential motor or ion channel proteins are likely coordinating the tonotopic sensory of sound along the cochlear axis.

### Spatiotemporal dynamics of communication between hair cells and SGNs reveal the tonotopic organization along cochlear axis

To understand how HCs cooperate with SGNs to sense and convey different-frequency sound signals, we interrogated secreted signaling events among cochlear epithelium and SGN for the three spatial datasets. At E17.5 stage, the maximum number of interactions is found between SGNs and HCs, when the SGNs were the hub (S6-1A and B Fig). This result was consistent with core function of SGNs for receiving the signals from HCs. Whereas, the stronger communications were observed between supporting cell subtypes and HCs when the HCs were the hubs (S6-1B, right). These results revealed that HCs were the center of cell connections in cochlea, communicating with nearby supporting cell types to coordinate its sensory function, not only just connecting with SGNs. At P8 stage, the number of interactions increased, either between SGNs and HCs or other cell types (Fig 6A, B). As for HCs, the strength of interaction with neighbor cells like inner sulcus cell and interdental cell (lateral Kolliker’s organ), as well as Hensen’s cell became stronger (Fig 6B and S6-1B Fig).

**Fig 6.**
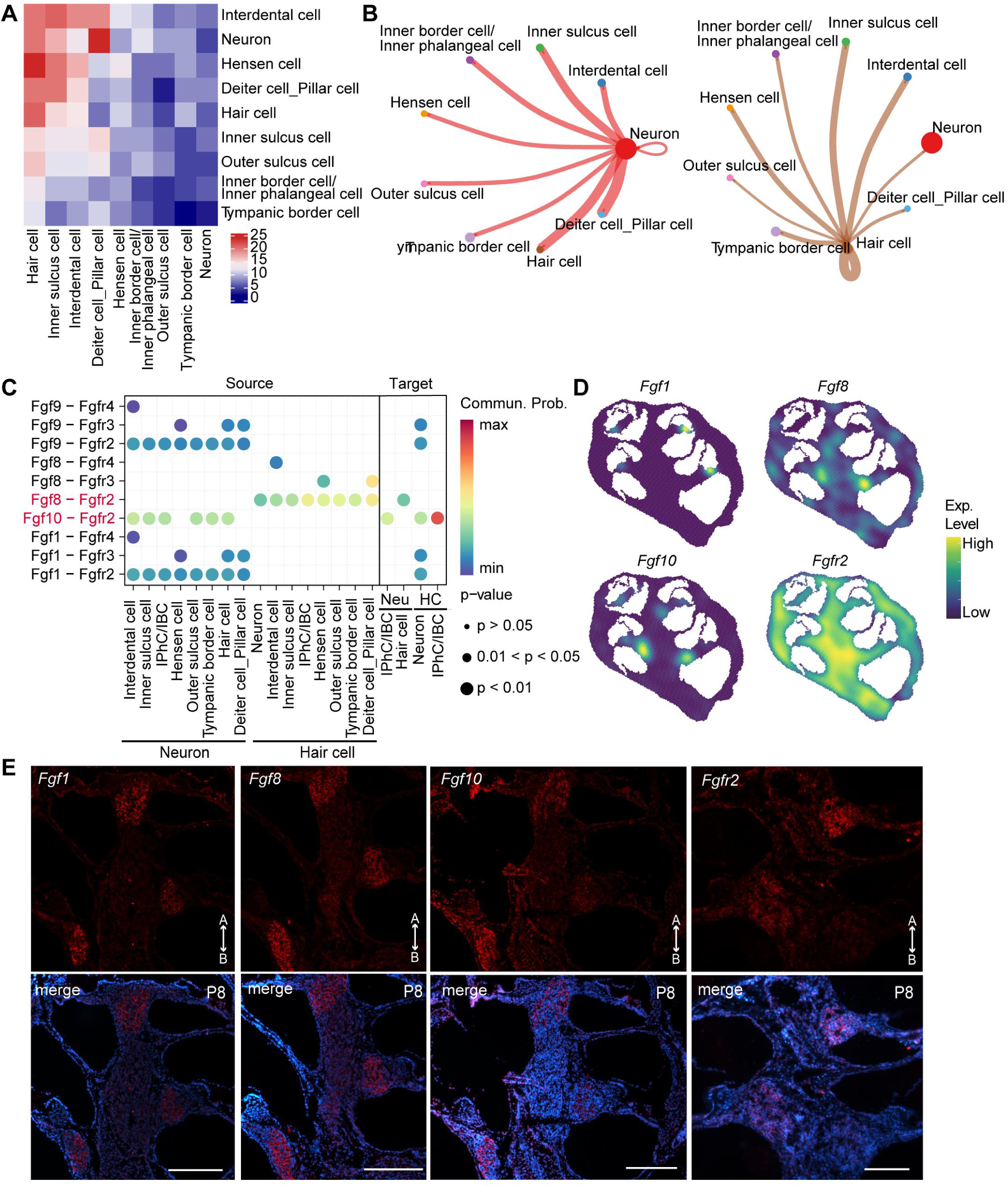
Developmental dynamics of cell-cell communication network between cell clusters from the organ of Corti. **(A)** Heatmap showing the number of ligand-receptor pairs evaluated by cellchat for neuron cell and cell clusters from organ of Corti (OC) at P8. **(B)** Left, number of significant ligand-receptor pairs between cell clusters from OC at P8, with the neuron cell as a hub. Right, number of significant ligand-receptor pairs between cell clusters from OC at P8, with hair cells as a hub. The edge width is proportional to the indicated number of ligand-receptor pairs. **(C)** Bubble plots showing ligand-receptor pairs of FGF signaling pathway between neuron cell and other cell clusters from OC. The size of dots represents the significance and the color represents the communication probability. **(D)** Spatial dense plots showing the expression level of *Fgf10*, *Fgfr2*, *Fgf1*, and *Fgf8* at P8 cochlea section. **(E)** RNA *in situ* validating the expression of *Fgf1*, *Fgf8*, *Fgf10*, and *Fgfr2* mRNA on P8 cochlea sections. Scale bar equals to 200 μm. A, apical; B, basal.

Next, all pathways were analyzed and ranked according to their communication intensity. The interaction strength of Ptn-Sdc, Mdk-sdc, and Fgf pathways were among the top 3 of all the interactions (S6-2 Fig). Fgf signaling pathway plays crucial roles in inner ear development and were further analyzed between these cell types of OC [42]. When neurons acted as the signaling source, it interacted with cells from the OC mainly by Fgf10-Fgfr2 signaling both at E17.5 and P8 stages (Fig 6C and S6-1C Fig). When HCs acted as the signaling source at P8, it mainly interacted with other cells from the OC by Fgf8-Fgfr2 signaling, especially with Deiter cell/pillar cell (Fig 6C). While Neurons as the signaling source mainly interacted with other cells by Fgf10-Fgfr2 (Fig 6C). With RNA *in situ* assay, the expression location of The *Fgf8* were confirmed, which were exclusively expressed in IHCs [43] and SGNs (Fig 6D and E). Strong expressions of *Fgf10* and *Fgfr2* were detected in OC and SGN regions (Fig 6D, E and S6-1 Fig). Interestingly, their expressions exhibited a gradient pattern in SGNs, with relatively stronger expression of *Fgf10* in the middle/basal SGNs and stronger *Fgfr2* in the apical SGNs (Fig 6D, E and S6-1, S6-2 Fig), suggesting spatial differences of pathway components to coordinate cell communications across the cochlear axis.

Next, regionalized cell communications were further analyzed across the cochlear axis. The OC cell types from apical, middle, and basal regions were extracted from the whole spatial single-cell atlas, and were subjected to differential cell communication studies. The overall cell communications were weak at the apical region, but relatively stronger in the middle and basal regions (Fig 7A-C). The strongest communications are found in the middle region (Fig 7B), which may serve as the hub regions in the cochlea. These regionalized heterogeneities of the connection intensity between SGNs and HCs (Fig 7A-C) were confirmed by the immunostaining assay with Tuj1 (also known as βIII-tubulin), which is a widely recognized marker for neuronal cells [44] and is often used for detecting the neuronal projections of SGNs in the inner ear [45, 46]. The immunostaining of Tuj1 revealed the strongest signal intensity of Tuj1 in the middle region rather than the apical and basal regions (Fig 7D, E), suggesting their strong connections in the middle region. These projection features of SGNs revealed by immunostaining (Fig 7A-C) were consistent with the differential communication studies in three different regions (Fig 7D, E). The counting of ligands and receptors along the apical-to-basal axis demonstrated that the Ptn-Sdc4 signaling pathway is the most dominant pathway in the apical SGNs or HCs, whereas the *Fgf10*-*Fgfr2* is the most dominant signaling hub in the basal SGNs (Fig 7-2 and S7-1, S7-3 Fig). This regional heterogeneity of cell signaling is likely dependent on the differential gradient expression of components of each signaling pathway (Fig 6 and S7-3 Fig), coordinating the tonotopic organization in the cochlea.

**Fig 7.**
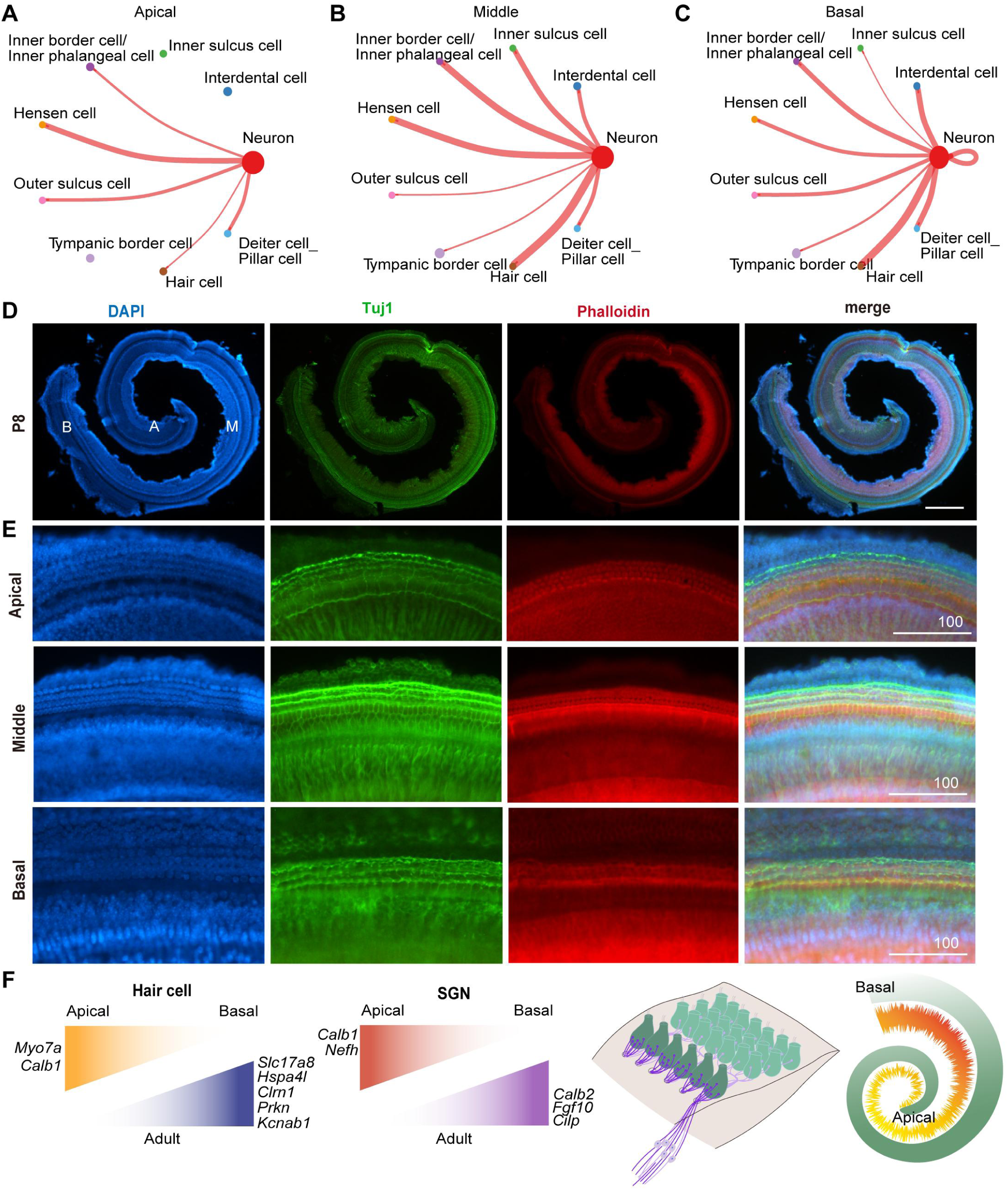
Spatial organization of hair cells, neuronal cells, and their communication network reveals the cochlear tonotopic organization. (A-C) Number of significant ligand-receptor pairs between neuron cell and other cell clusters from OC located in the apical, middle and basal regions at P8, respectively. The edge width is proportional to the indicated number of ligand-receptor pairs. **(D)** The whole-mount immunostaining of Tuj1 (green) and phalloidin (red) on P8 cochlea. DAPI, blue. Scale bar equals to 200 μm. **(E)** Magnified view of **(D)** to show neuronal connections and projections in apical (upper), middle (middle), and basal (bottom) regions, respectively. **(F)** Schematic representation illustrating the gradient patterning observed in adult hair cells and SGNs, highlighting a potential molecular mechanism underlying the tonotopic organization along the cochlear axis. A, apical; M, middle; B, basal.

## Discussion

In this study, the spatiotemporal transcriptomic analyses in cochlea revealed the developmental dynamics of HCs and SGNs and demonstrated their gradient gene expression patterns from apex-to-base, and their regionalized communications as well. These spatiotemporal data provided deeper insights into the molecular and cellular mechanisms underlying the tonotopic organization along cochlear axis. According to these results, a model was proposed to illustrate the underlying mechanism for tonotopic organization (Fig 7F and S7-1C Fig). The cochlea organizes its cell types in a spatially specific manner. The HCs and SGNs expressed a number of genes, like *Myo7a*/*Ocm* in OHCs, *vGlut3*/*Calb2* in IHCs, and *Calb1*/*Calb2*/*Nefh* in SGNs, in a gradient pattern from apex to basal, either in a decreased or increased manner, together finely regulating cochlear mechanical tuning (Fig 7F and S7-1C Fig). Accordingly, their communications are toned into a spatially regionalized pattern with divergent connection intensity from apical to basal regions. These divergent gradients patterns are mirrored by gradient features of sound frequency sensitivity along cochlear axis, which may be the molecular fundamentals for establishing the tonotopic organization (Fig 7F and S7-1C Fig).

We identified distinct spatial expression patterns of SGNs subtype marker genes, suggesting the spatial differentiation of SGN subtypes along the cochlear axis. Our previous work newly identified an additional SGN subtype, type III [33], which is very similar with type I SGNs according to their commonly expressed marker genes. Our results showed that the type III marker gene *Nefh* displayed a gradient pattern with high expression level predominantly in basal regions (Fig 3), suggesting that the type III SGNs dominantly located in the basal region. A recent study using conventional immunostaining demonstrated a differential distribution of SGN subtypes along the cochlear axis, with type Ia and Ib neurons primarily in apical and middle regions, while type Ic was more concentrated in the base [10]. However, the gene markers used in their study for distinguishing each SGN subtypes are different from ours, which may make certain inconsistent conclusions. They used Calb2 to label type Ia SGNs, Calb1 to label type Ib SGNs, and double negative Calb1^□^/Calb2^□^ to label type Ic SGNs [10]; while *Calb2* is dominantly expressed in type III and Ia according to our snRNA-seq data (S1-3B Fig). But the same conclusion is that the SGN subtypes exhibit a divergent distribution along apical-basal axis.

Interestingly, our results showed the gradient patterns of those marker genes for each SGN subtype were temporally dynamic, such as that the gradient expression of *Nefh* and *Calb2* in SGNs at embryonic stage was reversed in adulthood. These spatiotemporally dynamic patterns but similar molecular features among subtype Ia, Ib, Ic, and III suggested that each SGN subtype represented a cell state of type I SGNs across different cochlea regions or across different stages. Type III SGNs may represent a new subtype of type I SGNs [10]. The type III SGNs were not identified in previous studies likely due to different approaches (scRNA-seq or snRNA-seq) and different developmental stages (developmental or adult stages) used in these studies [13, 24, 26]. The loss of spatial information might be an additional reason. Traditional histological and immunostaining methods are limited in their ability to achieve comprehensive large-scale molecular characterization and differential expression analysis of SGN subtypes [10]. The close molecular characteristics of type I and III SGNs complicate their distinguishing. Our spatial data indicate that these subtypes may represent the same cell type (type I) exhibiting subtle molecular differences based on their spatial location along the cochlear axis, contributing to tonotopic organization [10, 33].

In addition to SGNs, HCs also exhibit spatial gradient heterogeneity along the cochlear axis. Our findings reveal notable differences between OHCs or IHCs located in the basal and apical regions. Classic markers like Myo7a show a pronounced gradient from apex to base in OHCs rather than IHCs, indicating distinct molecular identities. The IHC marker Slc17a8 displayed a similar gradient pattern in adult cochlea, with the weakest expression in the basal region (Fig 5H). Other markers, including Calb1 (S5-1A Fig), Kcnq4, BK channels, and Prestin, were shown in this study or were reported in previous publications to display gradient distributions in inner ear [9]. Motor proteins and ion channels are essential for sensory function [47] and are supposed to coordinate the tonotopic organization of the cochlea, playing crucial roles in frequency tuning. Despite some inconsistences due to limited hair cell numbers in our spatial dataset, many reported gradient genes were confirmed by our spatial data, confirming obvious spatial heterogeneity of HCs. Currently, no single-cell transcriptomic studies have reported the subtypes of OHCs and IHCs in mice due to very limited number of HCs captured. Our spatial data and immunostaining results provided clear evidences for the existence of distinct OHC and IHCS subtypes. Moreover, these findings suggested that HCs may also be spatiotemporally subdivided into distinct subtypes, very similar to the SGN subtypes. These HC or SGN subtypes, as marked by differential gradient expression of neural sensation related motors or ion channels, together regulate the frequency tuning of cochlear.

Importantly, our analysis uncovered numerous genes exhibiting gradient expression patterns along the cochlear axis, indicating that these gradients may contribute to the regional specialization or patterning of frequency sensitivity in the cochlea. Many gradient genes identified from earlier developing cochlea in previous publications were associated with the development [13, 24, 26]. They also constructed 3D spatiotemporal maps of the cochlea during the development, using scRNA-seq techniques [13]. However, it is likely that these gradient expression patterns were transient, subject to be reversed, or may have even disappeared in the later stages, based on our study. In contrast, the gradient genes identified in this study from the adult stage were primarily associated with energy metabolism and neural activity (Fig 5D). This association highlights the ultimate forms of cochlear organization and their sensory functions. Therefore, gradients from the adult stage, rather than the developmental stage, more likely align with the frequency discrimination patterns in the cochlea, highlighting the significance of the identifying gradient genes from the adult stage in this study. Furthermore, our data revealed regionalized differences in cell-cell communication within the cochlea, emphasizing the importance of regionalization in the inner ear’s functional organization. Understanding these localized interactions is crucial for uncovering the mechanisms governing cochlear development and its ability to perceive sound frequencies with high precision. Overall, this study highlights the temporal and regionalized gene expression patterns, along with the specialized organization and communication among cell types, including hair cells, SGNs, and supporting cells. This coordinated mechanism is essential for establishing tonotopic patterning during development and achieving precise sensory function.

Given the structural complexity of the inner ear, spatial organization is crucial for auditory function. Genetic variations disrupting this architecture often cause hearing defect [48, 49]. While single-cell RNA-seq has been widely used in inner ear development research and has yielded a wealth of insights into inner ear development, it lacks the crucial spatial context essential for understanding developmental and genetic mechanisms. In contrast, our spatial transcriptomic analysis preserves this spatial context, providing significant advantages for studying inner ear development and architecture. Our spatial transcriptomic analysis offers invaluable insights that address these challenges and will undoubtedly have significant implications for the field. We mapped the expression patterns of known deafness-associated genes on inner ear tissue, revealing spatial expression hotspots that identify structures most vulnerable to mutations causing hearing loss (S8 Fig). Many of gradient genes identified in this study are associated with deafness or hearing defects, such as *Myo7a*, *Calb2*, *Fgf10*, *Eya1*, and *Gata3*, whose mutation were reported to cause morphological abnormality of inner ear or hearing defects [50]. These spatial data offer important resources for the discovery of deafness-causing genes and investigation of the pathogenic mechanisms underlying hearing defects.

### Limitations of the study

Due to the significant value of the spatial transcriptomic data, our study has led to many interesting findings. However, there are still some challenges and limitations associated with the application of spatial transcriptomics technology in the context of the inner ear. The architecture of the cochlea is highly specialized and intricately organized, with each section differing from the others. A comprehensive analysis of all serial sections and 3D-construction may be required for a more complete characterization of inner ear architecture, as well as in combination with other cutting-edge technology, such as multiplexed error-robust *in situ* hybridization (MERFISH) [51]. Due to the inherent limitations of the spatial technology itself, spatial transcriptomics still has some intrinsic issues that are difficult to resolve [27, 52]. The spatial stacking between cells is unavoidable and the thickness of cell bodies left in a tissue section is also spatially dynamic, which can all compromise the resolution and accuracy of differential expressing gene studies. Moreover, we analyzed the spatial transcriptomic data at a resolution of 10 μm. This resolution may present challenges in accurate identification of densely packed cell types with small size. Even though our spatial transcriptomic data can be adjusted to a maximum resolution of 5 μm, this enhancement could simultaneously limit the number of detectable genes. Despite less resolution compared to previous inner ear scRNA-seq or snRNA-seq datasets, the spatial single-cell atlas established in this study is valuable for single-cell level transcriptomic analyses. The spatial datasets served as a valuable resource for researchers investigating inner ear development and auditory function, allowing for the exploration of spatiotemporal gene expression patterns of any gene of interest, or any particular region of interest.

## Materials and methods

### Experimental model

Wild-type C57BL/6 mice were bred in the laboratory. All animal experiments were conducted with approval from the Animal Research and Ethics Committee of the Chengdu Institute of Biology, Chinese Academy of Sciences (Approval No. CIBDWLL2018030). Mice were provided clean bedding, ad libitum water and food, and nestlets with 12 h light/dark cycles in ventilated cages. Sex of mouse samples in this study was not characterized.

### Tissue collection

For E17.5 cochlear tissue, a pregnant female was euthanized and inner ears were dissected from pups. For postnatal time points, P8 and adult mouse (two-month-old) were euthanized and inner ears were dissected. Cochlea were separated from the inner ears under the stereo microscope (Olympus, SZ61) and washed by ice-cold DPBS. After wiping the liquid by dust-free paper, cochleae were rinsed with ice-cold OCT, and next moved into Cassettes with ice-cold OCT. In order to gain better longitudinal section with at least 4 cochlea duct regions, the tissue position in the cassette was fine-tuned under the stereo microscope again. Then the cassettes were quickly put on the dry ice to ensure the tissues were froze immediately. All samples were sent to Beijing Biomarker Technologies Co. the same day.

### Tissue cryosection and processing for spatial-seq

The spatial-seq was performed with the BMKMANU S1000 Gene Expression kit (BMKMANU, ST03002). Tissue sectioning, RNA quality, HE staining, imaging and tissue optimization were followed by the BMKMANU S1000 Tissue Optimization kits user guide (BMKMANU, ST03003). The Cryosections were cut at a thickness of 10 µm and every section would be checked with HE staining before target structure section appearing. When 4 cochlea duct holes were observed, the next section would be moved to slide for later sequencing.

### Library preparation and sequencing

Library constructions were also performed according to the BMKMANU S1000 Library Construction Kit User Guide. The Illumina library was sequenced with Illumina NavoSeq 6000. The raw data from Illumina and Nanopore were mapped to the mouse reference genome (GRCm38, ftp://ftp.ensembl.org/pub/release-74/fasta/mus_musculus/) by BSTMatrix v2.4 (http://www.bmkmanu.com/wp-content/uploads/2024/07/BSTMatrix_v2.4.f.1.zip) and BSTMatrix-ONT v1.2 (http://www.bmkmanu.com/wp-content/uploads/2023/03/BSTMatrix-ONT_v1.2.zip), respectively, using default parameters. The image adjusted by BSTViewer V4.7 (http://www.bmkmanu.com/wp-content/uploads/2024/07/BSTViewer-V4.7.4.1.rar) and corresponding level 2 (10 μm) matrix were used for downstream analysis. A total of 5,545, 10,214 and 7,414 spots were obtained from single E17.5, P8 and 2M mouse cochlea section samples at a resolution of 10 μm, respectively.

### Cell clustering and cell-type identification

Downstream analysis of BSTViewer output data was performed using the R package Seurat (v5.1.0) (https://github.com/satijalab/seurat) [53]. HE images from the same sample were pooled together as a data set for analysis. First, for all samples, data was preliminary filtered with min.cell at 5 and min.fearture at 100, when imported into R. Then all datasets were normalized (scale.factor = 10000) and scaled using all genes. Principal component analysis (PCA) focusing on the most variable features was performed and for all data, the first 30 dimensions obtained by RunPCA were applied to the analysis of RunUMAP, FindNeighbors, and FindClusters to obtain UMAP and cluster information. Cluster specific genes were identified by FindAllMarkers differential expression analysis. Combining cell-type specific markers identified by Seurat and previously reported markers, spot’ ident was annotated.

### Cell communication analysis of hair cells and supporting cells

Cellchat (v1.6.1)[54] package was applied to explore the potential interaction pattern of cell types from OC and SGN. Based on the normalized gene-expression matrix of our datasets, the database of mouse ligand-receptor interactions was used to identify overexpressed genes and interactions with the method of “trimean”, compute the communications and related pathways. All analysis steps were followed by the suggested workflow of CellChat with the default parameters.

### Pseudotime trajectory analysis of HC and SGN groups

The newCellDataSet function of Monocle2 (v2.32.0) [55] was utilized to create a CellDataSet object, and the estimateSizeFactors and estimateDispersions functions were applied for the estimation of size factors and dispersions, respectively. By employing the detectGenes function, genes expressed in at least 10 cells were identified, and key genes for trajectory inference were set using the setOrderingFilter function. Subsequently, the reduceDimension and orderCells functions were applied for dimensionality reduction and cell ordering to construct the pseudotime trajectory of cells. The plot_cell_trajectory function was used to visualize the cellular trajectories, which were colored according to cell states. Additionally, genes with expression changes during cell development and genes that regulate the branching points of cellular differentiation were identified using the differentialGeneTest and BEAM functions. Finally, a heatmap of pseudotime-associated genes was generated using the plot_pseudotime_heatmap function to display the expression patterns of these genes across different cellular states.

### Gene expression pattern analysis from apex to basal part

To explore regional gene-expression characteristics along the cochlea duct, the expression matrixes of SGN and organ of Corti cell types from apex, middle-basal and basal reagions (Fig. 4a; Fig S4a and d) were extracted separately for DEG analysis, with the method of wilcoxon test. To better reveal the expression pattern of differential genes, plot_density method from Nebulosa (v1.14) package[56] was used to depict spatial expression.

### Immunofluorescence

Cochleae from E17.5, P8 and Adult mice were fixed in 4% paraformaldehyde diluted in 1×PBS for 2 hours at 4 [, washed in 1×PBS for three times, dehydrated by 30% sucrose solution, then embedded in OCT Compound for cryosectioning at 10 μm thickness. According to previous publications [43], the immunofluorescence was performed as following steps. Briefly, sections were blocked in 5% bovine serum albumin (BSA, Biofroxx, 1110GR500)/10% horse serum albumin (HSA, Sangon Biotech, E510006-0100) / 1% Triton-X/1×PBS for 8∼10 hours at 4 [ and primary antibodies were applied overnight at 4 □ in 5% BSA/10% HSA/0.5% Triton-X/1×PBS. Secondary antibodies were applied in 5% BSA/10% HSA/0.5% Triton-X/1×PBS for 6∼8 hours at 4 □ followed by DAPI nuclear staining. All washes following primary and secondary antibody application were performed with 1% Triton-X/1×PBS. All fluorescent images were acquired using fluorescent microscope system (Nikon, Eclipse 55i). All antibodies used in this study were listed in S4 File.

### RNA *in situ* hybridization

The fragments of target genes (S4 File) were prepared by PCR amplification of cDNA from cochleae RNA, with reverse primers containing a T7 RNA polymerase promoter sequence. Next the DIG-labeled RNA probes were synthesized by the transcription reaction using Riboprobe Systerm-T7 kit (Promega, P1440) and DIG labeled UTP (Roche, 11277073910). The slides for RNA *in situ* hybridization (RNA-ISH) were prepared with the same steps of immunofluorescence. Then RNA-ISH was conducted as following steps, according to previous publications [33, 57]. Firstly, slides were fixed by 4% PFA for 5 min at RT and washed in 1×PBS for 3 min and in distilled water for 1 min. Next for permeabilizing, slides were immersed in 0.2 M HCl for 15 min and 0.1 M triethanolamine-HCl for 10 min. After being washed in 1×PBS for 5 min, slides were dehydrated by washing for 20 min per wash in 75%, 85% and 100% ethanol. And then slides were treated by hybridization buffer containing DIG-labeled RNA probes overnight at 65 □ To remove non-specific and repetitive RNA hybridization, slides were washed in 5×saline sodium citrate (SSC) for 5 min at RT and 2×SSC for 30 min at 60 □ After being treated in 0.3% H_2_O_2_ for 1 h at RT, slides were blocked for 2 h at RT and next incubated with anti-DIG antibody overnight at 4 □ Finally, the signal was detected by the TSA® Plus Cyanine 3.5 (Cy3.5) detection kit (Akoya, NEL763001KT) and slides were sealed by diluted DAPI with glycerin. The images were captured by fluorescent microscope.

### Quantificatin and statistical analysis

All statistical analyses were conducted by R and RStudio. And as for the results of wilcoxon-test, which were used to uncover the spatial differential expression genes for neuron cells and OHCs, *P*-value < 0.05 were considered statistically significant.

## Acknowledgments

The work was supported by the National Natural Science Foundation of China (Nos. 82300142, 82301316, 32370882, and 32370536), and the Central Government Guides Local Science and Technology Development Funds in Sichuan Provincial (2022ZYD0131), and the Fundamental Research Funds for the Central Universities (BMU2021YJ064). We gratefully acknowledge the High-performance Computing Platform of Peking University for conducting the bioinformatics analysis.

## Author contributions

J. Li, Y. Zhao, and H. Wu designed the work; M. Yan, P. Zhang, H. Wang, J.H. Li, Y. Wang, X. Zeng, and Y. Gao acquired the data; M. Yan, H. Wang, Y. Wang, B. Zhu, D. Deng, F. Deng, X. Guo, L. Ma and Y. Feng analyzed the data; M. Yan, Y. Wang, and H. Wang made the figures; J. Li, Y. Zhao, and H. Wu wrote the paper.

## Supporting information

**S1-1 Fig. Spatial single-cell atlas of P8 cochlea. (A)** Bubble plot analysis of the top differentially expressed genes for each cell types in the P7 cochlea (left, scRNA data from reported assay) and P8 cochlea (right, spatial scRNA data). The size of the dots represents the percentage of cells in the clusters expressing the gene and the color intensity represents the average expression levels of the gene in that cluster. **(B)** Density plots showing the spatial expression pattern of several cell-type markers in the P8 cochlea section chip.

**S1-2 Fig. Spatial single-cell atlas of E17.5 cochlea. (A)** Spatial single cell types on the E17.5 cochlea cross-section. Seven major groups of cell types were shown, respectively. **(B)** Bubble plot analysis of the top differentially expressed genes for each cell types in the E16 cochlea (left, scRNA data from reported assay) and E17.5 cochlea (right, spatial scRNA data). The size of the dots represents the percentage of cells in the clusters expressing the gene and the color intensity represents the average expression levels of the gene in that cluster. **(C)** Correlation heatmap showing the correspondence of cell annotation between spatial cells from E17.5 cochlea and the reported scRNA-seq dataset from E16 cochlea. OS: Out structure, EryC: Erythrocytes, MaC: Macrophages, EnC: Endothelial cell, NeC: Neutrophils cell, FC: Spiral ligament fibrocyte, SLg_FC: Spiral ligament fibrocyte, SLb_FC: Spiral limbus fibrocyte, TBC: Tympanic border cell, Sch: Schwann cell, RMC: Reissner’s membrane cell, BS: Basal stria, MS: Marginal stria, IS: Intermediate stria, IdC: Interdental cell, KO:, Kölliker’s organ cell, CC/OSC: Claudius cell/Outer sulcus cell, SGN: Spiral ganglion neuron, HC: Hair cell, DC/PC: Deiter cell/Pillar cell, SV: Stria vascularis, IdC/ISC: Interdental cell/Inner sulcus cell, L.KO: Lateral kölliker’s organ cell, IPhC: Inner phalangeal cell, HeC: Hensen cell, LER: Lesser epithelial ridge cell, IPC: Inner pillar cell. **(D)** Density plots showing the spatial expression pattern of several cell-type markers in the E17.5 cochlea cross section.

**S1-3 Fig. Spatial single-cell atlas of adult cochlea. (A)** Spatial single cell types on the adult cochlea cross-section. Eight major groups of cell types were shown, respectively. **(B)** Bubble plot analysis of the top differentially expressed genes for each cell types in the adult cochlea (left, scRNA data from reported assay; right, spatial scRNA data). The size of the dots represents the percentage of cells in the clusters expressing the gene and the color intensity represents the average expression levels of the gene in that cluster. **(C)** Correlation heatmap showing the correspondence of cell annotation between spatial dataset and the reported scRNA-seq dataset for adult cochlea. PeC: Pericytes, MaC: Macrophages, EnC: Endothelial cell, FC1: Fibrocyte 1, FC1: Fibrocyte 2, Ia_SGNs: Type Ia Spiral ganglion neuron, Ib_SGNs: Type Ib Spiral ganglion neuron, Ic_SGNs: Type Ic Spiral ganglion neuron, II_SGNs: Type II Spiral ganglion neuron, III_SGNs: Type III Spiral ganglion neuron, I_Sch: Type I schwann cell, II_Sch: Type II schwann cell, III_Sch: Type III schwann cell, RMC: Reissner’s membrane cell, BS: Basal stria, MS: Marginal stria, IS: Intermediate stria, CC/OSC: Claudius cell/Outer sulcus cell, IPhC/HeC: Inner phalangeal cell/Hensen cell, TBC: Tympanic border cell, OHC: Outer hair cell, IHC: Inner hair cell, PC: Pillar cell, OS:, NeC:, OsC:, SLg_FC: Spiral ligament fibrocyte, SLb_FC: Spiral limbus fibrocyte, SGN: Spiral ganglion neuron, SchC: Schwann cell, IdC: Interdental cell, DC/PC: Deiter cell/Pillar cell. **(D)** Density plots showing the spatial expression pattern of several cell-type markers in the adult cochlea cross section.

**S2 Fig. Psudotime analysis of developing spiral ganglion neurons. (A)** Plots showing different marker genes delineating different-stage neuron trajectories. **(B)** Heatmap showing dynamic expression changes of genes. **(C)** Related GO terms of genes from neurons at different clusters in **(B)**. **(D)** Curve plots showing the expression changes of the functional genes in developing neurons along pseudotime.

**S3-1 Fig. Spatial differential analyses of neuronal expressing genes from apex to base at E17.5 and adult stages. (A)** Upper left, spatial distribution of neuron cells at E17.5 cochlea cross section, divided by rectangular boxes with different colors (three regions: A, apex; M, middle; B, base). Bottom left, UMAP plot showing the apical, middle and basal neurons extracted from E17.5 cochlea as shown on the left, as indicated by different colors. Right, heatmap showing the expression level of the gradient genes representing two patterns in E17.5 SGNs (decreasing or increasing expression pattern from apical to basal regions). **(B)** Upper left, spatial distribution of neuron cells at adult cochlea cross section, divided by rectangular boxes with different colors (three regions: A, apex; M, middle; B, base). Bottom left, UMAP plot showing the apical, middle and basal neurons extracted from adult cochlea as shown on the left, as indicated by different colors. Right, heatmap showing the expression level of the gradient genes representing two patterns in adult SGNs (decreasing or increasing expression pattern from apical to basal regions). **(C)** Volcano plots showing the differential analysis of spatial expression in SGNs from E17.5, P8 and adult cochlea. (**D**) Density plots showing the spatial expression patterns in E17.5 and adult SGNs on cochlea cross section, representing two gradient patterns (apex-to-base decreasing and increasing expression). **(E, F)** The whole amount view showing the immunostaining of Calb1 **(F)** and Calb2 **(G)** in adult cochlea. A, apex; M, middle; B, base. (**H**) Spatial expression and immunostaining of Pou4f1 in adult cochlea cross section. **(G)** Gene ontology analysis of spatial gradient genes from E17.5 and adult SGNs.

**S3-2 Fig. Spatial dynamics of spiral ganglion neurons subtypes along the cochlear axis. (A**-**E)** Immunostaining of Calb1 **(A)**, Calb2 **(B)**, Pou4f1**(D)** and RNA in situ detection of *Cilp* mRNA **(C)** on the adult cochlea cross-sections. DAPI is labeled in blue. A, apex; B, base. Scale bar equals to 200 μm.

**S4 Fig. Spatiotemporal expression changes of hair cell markers. (A, B, C)** Whole-mount immunostaining of Myo7a in cochlear hair cells at E17.5 **(A)**, P8 **(B)** and adult **(C)**. Myo7a is labeled in green; phalloidin, red; DAPI, blue. Scale bars equal to 200 μm. (**D**) Magnified view of (**C**) showing apical (upper), middle (middle), and basal (bottom) regions of adult cochlea, respectively. A, apical; M, middle; B, basal. Scale bars equal to 100 μm.

**S5-1 Fig. Spatial differential analysis of hair cell markers from apex to base at adult. (A)** Whole-mount immunostaining and magnified view of Calb1 (green) in the cochlear hair cells. (**B, C**) Whole-mount immunostaining and magnified view of Foxp1 (green) in the adult cochlea. DAPI is labeled in blue. Scale bars equal to 200 μm.

**S5-2 Fig. Spatial differential analysis of OC from apex to base at E17.5 and P8 stages. (A)** Upper, spatial distribution showing cell clusters from OC identified in E17.5 and adult cochlea cross section, divided by rectangular boxes with different colors (three regions: A, apex; M, middle; B, base). Bottom, heatmap showing the expression level of the gradient genes representing two patterns in E17.5 and P8 OC (decreasing or increasing expression pattern from apical to basal regions), respectively. **(B)** Density plots showing the spatial expression patterns in E17.5 and adult OC on cochlea cross section, representing two gradient patterns (apex-to-base decreasing and increasing expression). **(C)** Bar plot showing GO analysis of spatial gradient genes from E17.5 and adult OC. (**D**) Bubble plot showing the expression levels of a set of genes representing two gradient patterns (apex-to-base decreasing and increasing expression) in OHCs on P8 cochlea sections. (**E**) Network plot showing several significant GO terms and its related genes enriched in the apical OC of E17.5 and P8 cochlea, respectively.

**S5-3 Fig. Spatial differential analysis of HCs from apex to base at E17.5 and P8.** (**A**) UMAP plot showing HCs extracted from apical, middle and basal regions of E17.5 cochlea. (**B**) Bubble plot showing the expression levels of a set of genes representing two gradient patterns (apex-to-base decreasing and increasing expression) in HCs from E17.5 cochlea. The size of the dots represents the percentage of cells in the clusters expressing the genes and the color intensity represents the average expression levels of the gene in that cluster. (**C**) UMAP plot showing HCs extracted from apical, middle and basal regions of P8 cochlea. (**D**) Bubble plot showing the expression levels of a set of genes representing two gradient patterns (apex-to-base decreasing and increasing expression) in HCs from P8 cochlea. (**E**) Immunostaining of Slc26a5 (green), and phalloidin (red) in the P8 whole amount cochlea. DAPI is labeled in blue. A, apical; M, middle; B, basal. Scale bars equal to 200 μm. (**F**) Magnified view of (**E**) to show the variant expression level of Slc26a5 in HCs from apical (upper), middle (middle) and basal (bottom) regions, respectively. Scale bars equal to 100 μm.

**S6-1 Fig. Cell-cell communication network among cell clusters from the organ of Corti. (A)** Heatmap showing the number of ligand-receptor pairs evaluated by cellchat for neuron cell and cell clusters from OC at E17.5 **(B)** Left, number of significant ligand-receptor pairs of neuron cells (hub) interacted with cell clusters from OC at E17.5. Right, number of significant ligand-receptor pairs between hair cells (hub) and neuron cells as well as other cell clusters from OC at E17.5. The edge width is proportional to the indicated number of ligand-receptor pairs. **(C)** Bubble plots showing ligand-receptor pairs of FGF signaling pathway between neuron cells and cell clusters from OC at E17.5. The size of dots represents the significance and the color represents the communication probability. **(D)** RNA *in situ* detection of *Fgf10* and *Fgfr2* mRNA in the apical region of P8 cochlea sections.

**S6-2 Fig. Cell-cell communication network among neuron cells and cell clusters from the organ of Corti.** Bubble plots showing ligand-receptor pairs between neuron cells and other cell clusters from the organ of Corti for the P8 spatial data. The size of dots represents the significance and the color represents the communication probability.

**S7-1 Fig. Spatial organization of hair cells, neuronal cells, and their communication network reveals the cochlear tonotopic organization. (A, B)** Bubble plots showing ligand-receptor pairs of signaling pathway in the apical **(A)** and basal SGNs **(B)** at P8 stage. The size of dots represents the significance and the color represents the communication probability. **(C)** Schematic representation illustrating the gradient patterning observed in hair cells and SGNs (adult and P8 stage), highlighting a potential molecular mechanism underlying the tonotopic organization along the cochlear axis.

**S7-2 Fig. Spatial cell-cell communication network in hair cells. (A, B)** Bubble plots showing ligand-receptor pairs of signaling pathways which apical **(A)** and basal **(B)** HCs sent to other cell types in the organ of Corti (OC), respectively. **(C, D)** Bubble plots showing ligand-receptor pairs of signaling pathways which apical **(C)** and basal **(D)** HCs received from other cell types in OC, respectively. The size of dots represents the significance and the color represents the communication probability.

**S7-3 Fig. Spatial cell-cell communication network among neuron cells and cell clusters from the organ of Corti. (A-C)** Number of significant ligand-receptor pairs between neuron cells and cell clusters from the organ of Corti (OC) distributed in the apical **(A)**, middle **(B)**, and basal **(C)** regions at E17.5 stage, respectively. Up, the neurons are the hub; bottom, the hair cells are the hub. The edge width is proportional to the indicated number of ligand-receptor pairs. **(D, E)** Stack diagram comparing the weight **(D)** and number **(E)** of interactions every signaling pathway predicted between neuron cells and cell clusters from OC distributed in the apical, middle and basal regions at E17.5 stage, respectively. **(F)** Bubble plots showing that neuron cells and hair cells are the source sending information to other cell clusters from OC in the different regions at E17.5 stage. The size of dots represents the significance and the color represents the communication probability. **(G)** Bubble plots showing the ligand-receptor pairs that neuron cells and hair cells are the targets to receive information from other cell clusters from OC in the different regions at E17.5 stage. The size of dots represents the significance and the color represents the communication probability.

**S8 Fig. Spatiotemporal expression pattern of known deafness genes (A)** Density plots showing the spatial expression of all known deafness genes, on E17.5, P8 and Adult cochlea section. Strong densities were found at the organ of Corti (OC) region on E17.5 section, at OC and SGN region on P8 section, and at the central modiolus region on adult section. **(B)** Hierarchical clustering of cochlear cell types and deafness genes in spatiotemporal transcriptomic data. OC: the organ of Corti, FB: fibrocytes, LW: lateral wall, SGNs: spiral ganglion neurons, RM: reissner’s membrane, SC: Schwann cell, SS: surrounding structures.

**S1 File. All celltype markers of the three datasets, related to Fig 1 and S1 Fig.**

**S2 File. Gradient expression genes of Neurons in the three datasets, related to Fig 3 and S3 Fig.**

**S3 File. Gradient expression genes of HCs in the three datasets, related to Fig 5 and S5 Fig.**

**S4 File. Primers for *in situ* hybridization and antibodies for immunofluorescence.**

## Notes

https://ngdc.cncb.ac.cn/gsa

### Competing Interest Statement

The authors have declared no competing interest.

### Summary of Updates

author affiliations updated;Figure 1-5 updated;Supplemental files updated

