## Supplementary material for "Spatiotemporal transcriptomic analyses reveal molecular gradient patterning during development and the tonotopic organization along the cochlear axis": Supplymentary figures

#### S1-1 Fig

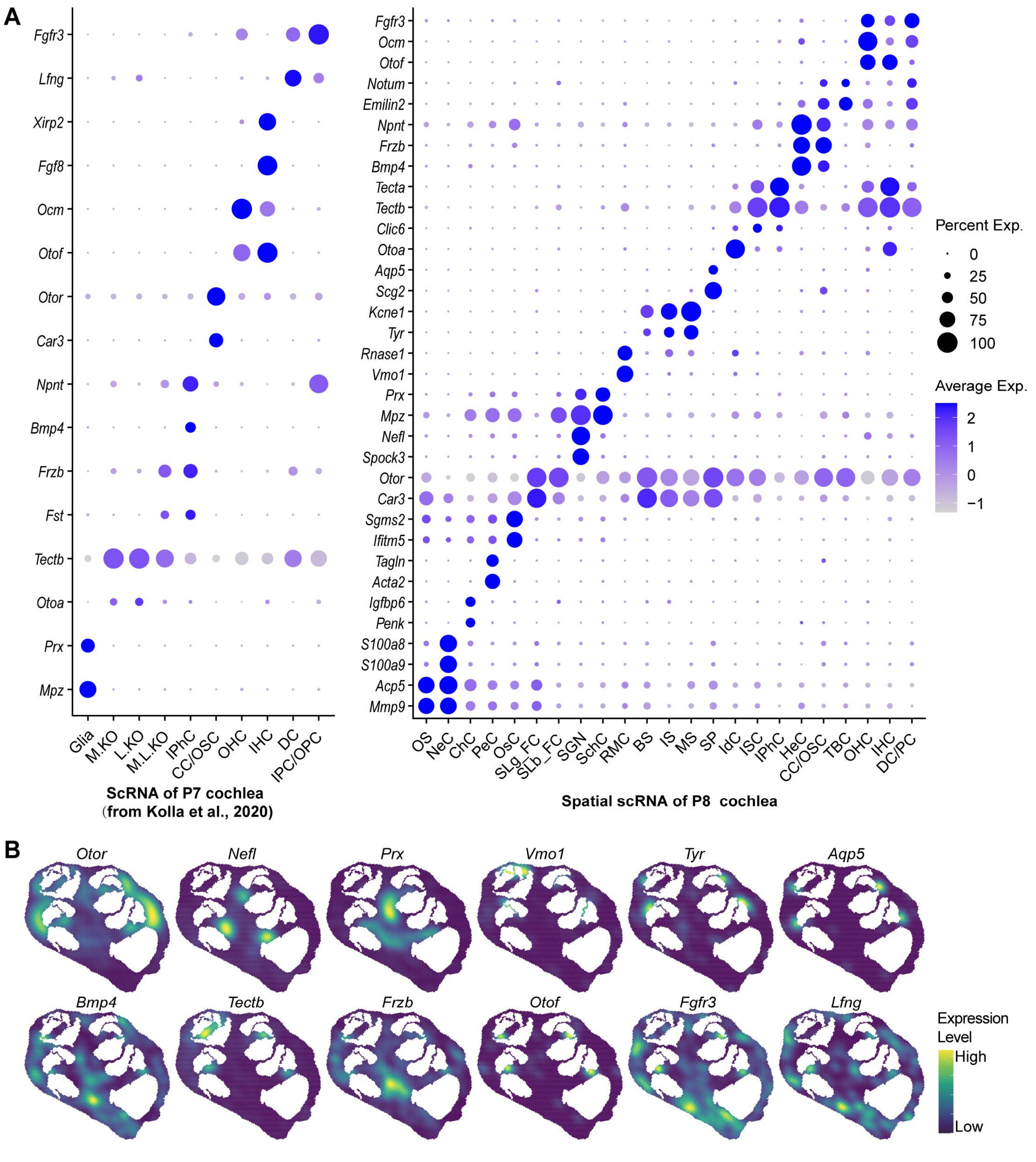
A
Fgfr3
Lfng
Xirp2
Fgf8
Ocm
Otof
Otor
Car3
Npnt
Bmp4
Frzb
Fst
Tectb
Otoa
Prx
Mpz
Glia
M.KO
L.KO
M.L.KO
IPhC
CC/OSC
OHC
IHC
DC
IPC/OPC
ScRNA of P7 cochlea
(from Kolla et al., 2020)
Fgfr3
Ocm
Otof
Notum
Emilin2
Npnt
Frzb
Bmp4
Tecta
Tectb
Clic6
Otoa
Aqp5
Scg2
Kcne1
Tyr
Rnase1
Vmo1
Prx
Mpz
Nefl
Spock3
Otor
Car3
Sgms2
Ifitm5
Tagln
Acta2
Igfbp6
Penk
S100a8
S100a9
Acp5
Mmp9
OS
NeC
ChC
PeC
OsC
SLg_FC
SLb_FC
SGN
SchC
RMC
BS
IS
MS
SP
IdC
ISC
IPhC
HeC
CC/OSC
TBC
OHC
IHC
DC/PC
Spatial scRNA of P8 cochlea
Percent Exp.
0
25
50
75
100
Average Exp.
2
1
0
−1
B
Otor
Nefl
Prx
Vmo1
Tyr
Aqp5
Bmp4
Tectb
Frzb
Otof
Fgfr3
Lfng
Expression Level
High
Low

S1-2 Fig

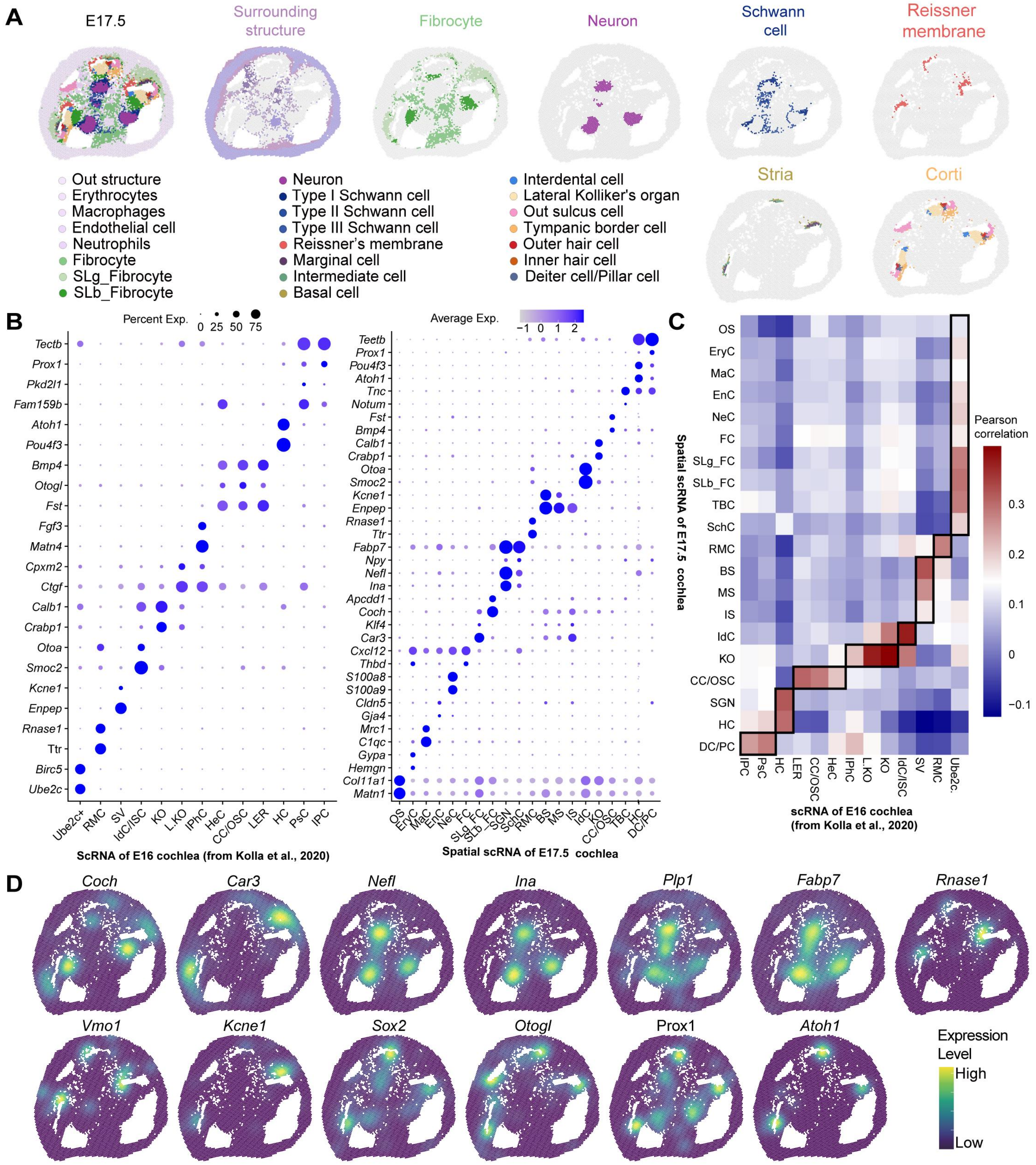
A
E17.5
Surrounding structure
Fibrocyte
Neuron
Schwann cell
Reissner membrane
Stria
Corti
Out structure
Erythrocytes
Macrophages
Endothelial cell
Neutrophils
Fibrocyte
SLg_Fibrocyte
SLb_Fibrocyte
Neuron
Type I Schwann cell
Type II Schwann cell
Type III Schwann cell
Reissner's membrane
Marginal cell
Intermediate cell
Basal cell
Interdental cell
Lateral Kolliker's organ
Out sulcus cell
Tympanic border cell
Outer hair cell
Inner hair cell
Deiter cell/Pillar cell
B
Percent Exp.
0 25 50 75
Average Exp.
−1 0 1 2
ScRNA of E16 cochlea (from Kolla et al., 2020)
Spatial scRNA of E17.5 cochlea
C
Pearson correlation
Spatial scRNA of E17.5 cochlea
scRNA of E16 cochlea (from Kolla et al., 2020)
D
Coch
Car3
Nefl
Ina
Plp1
Fabp7
Rnase1
Vmo1
Kcne1
Sox2
Otogl
Prox1
Atoh1
Expression Level
High
Low

### S1-3 Fig

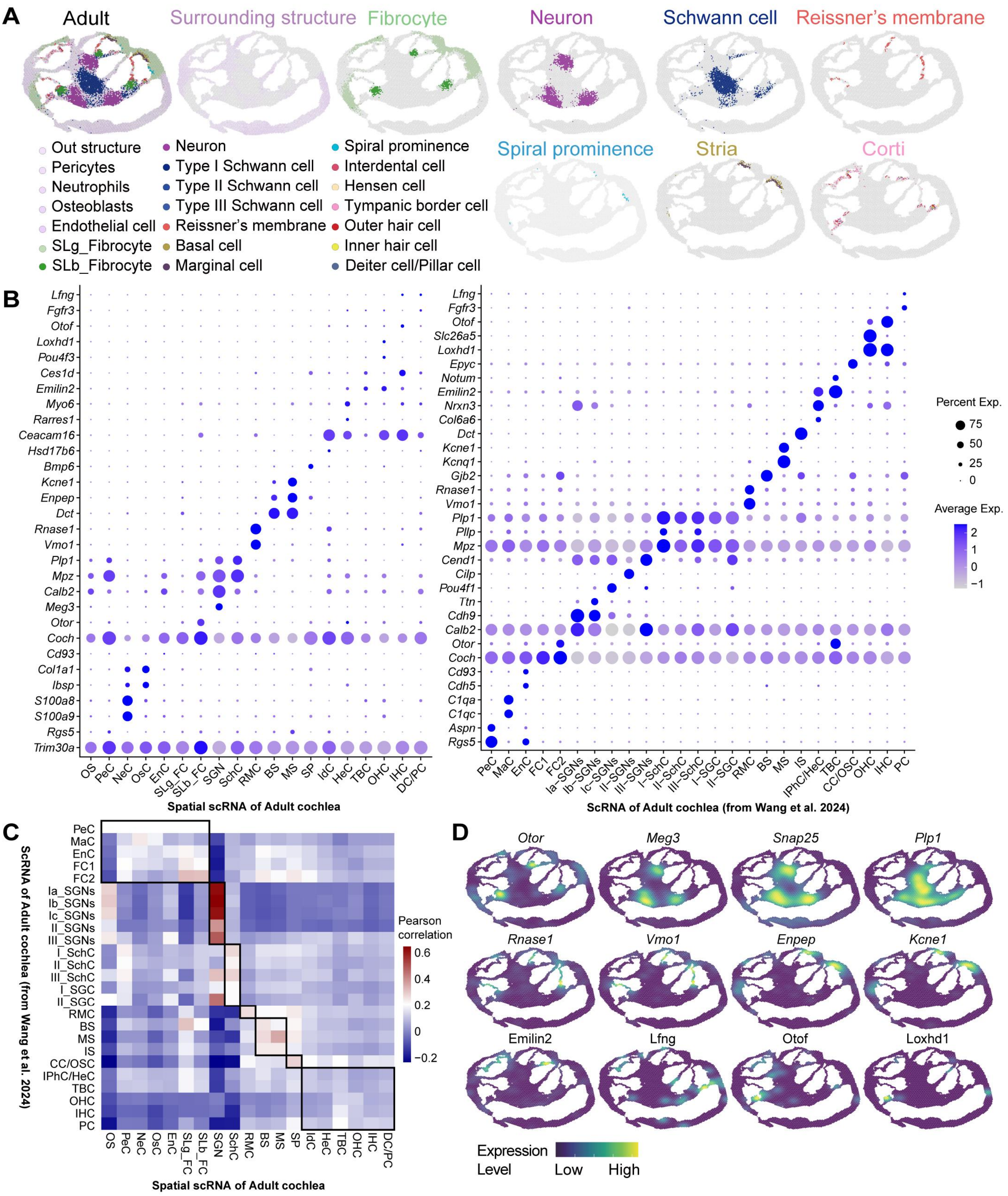
A
Adult
Surrounding structure
Fibrocyte
Neuron
Schwann cell
Reissner's membrane
Spiral prominence
Stria
Corti
Out structure
Pericytes
Neutrophils
Osteoblasts
Endothelial cell
SLg_Fibrocyte
SLb_Fibrocyte
Neuron
Type I Schwann cell
Type II Schwann cell
Type III Schwann cell
Reissner's membrane
Basal cell
Marginal cell
Spiral prominence
Interdental cell
Hensen cell
Tympanic border cell
Outer hair cell
Inner hair cell
Deiter cell/Pillar cell
B
Spatial scRNA of Adult cochlea
ScRNA of Adult cochlea (from Wang et al. 2024)
Percent Exp.
Average Exp.
C
ScRNA of Adult cochlea (from Wang et al. 2024)
Spatial scRNA of Adult cochlea
Pearson correlation
D
Otor
Meg3
Snap25
Plp1
Rnase1
Vmo1
Enpep
Kcne1
Emilin2
Lfng
Otof
Loxhd1
Expression Level
Low
High

#### S2 Fig

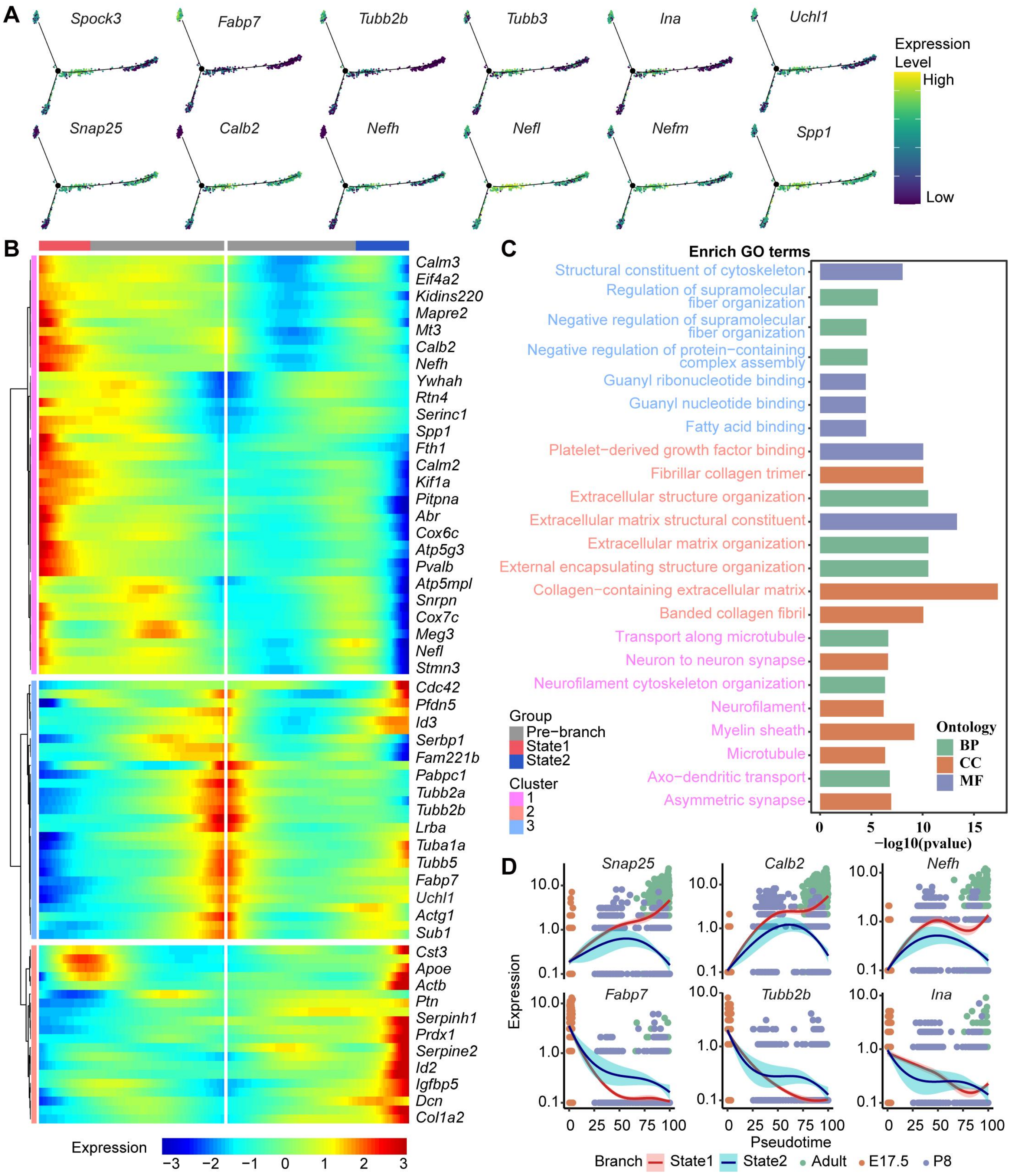
A
Spock3
Fabp7
Tubb2b
Tubb3
Ina
Uchl1
Snap25
Calb2
Nefh
Nefl
Nefm
Spp1
Expression Level
High
Low
B
Calm3
Eif4a2
Kidins220
Mapre2
Mt3
Calb2
Nefh
Ywhah
Rtn4
Serinc1
Spp1
Fth1
Calm2
Kif1a
Pitpna
Abr
Cox6c
Atp5g3
Pvalb
Atp5mpl
Snrpn
Cox7c
Meg3
Nefl
Stmn3
Cdc42
Pfdn5
Id3
Serbp1
Fam221b
Pabpc1
Tubb2a
Tubb2b
Lrba
Tuba1a
Tubb5
Fabp7
Uchl1
Actg1
Sub1
Cst3
Apoe
Actb
Ptn
Serpinh1
Prdx1
Serpine2
Id2
Igfbp5
Dcn
Col1a2
Expression
−3 −2 −1 0 1 2 3
Group
Pre−branch
State1
State2
Cluster
1
2
3
C
Enrich GO terms
Structural constituent of cytoskeleton
Regulation of supramolecular fiber organization
Negative regulation of supramolecular fiber organization
Negative regulation of protein−containing complex assembly
Guanyl ribonucleotide binding
Guanyl nucleotide binding
Fatty acid binding
Platelet−derived growth factor binding
Fibrillar collagen trimer
Extracellular structure organization
Extracellular matrix structural constituent
Extracellular matrix organization
External encapsulating structure organization
Collagen−containing extracellular matrix
Banded collagen fibril
Transport along microtubule
Neuron to neuron synapse
Neurofilament cytoskeleton organization
Neurofilament
Myelin sheath
Microtubule
Axo−dendritic transport
Asymmetric synapse
Ontology
BP
CC
MF
0 5 10 15
−log10(pvalue)
D
Snap25
Calb2
Nefh
Fabp7
Tubb2b
Ina
Expression
10.0
1.0
0.1
0 25 50 75 100
Pseudotime
Branch State1 State2 Adult E17.5 P8

**S3-1 Fig**

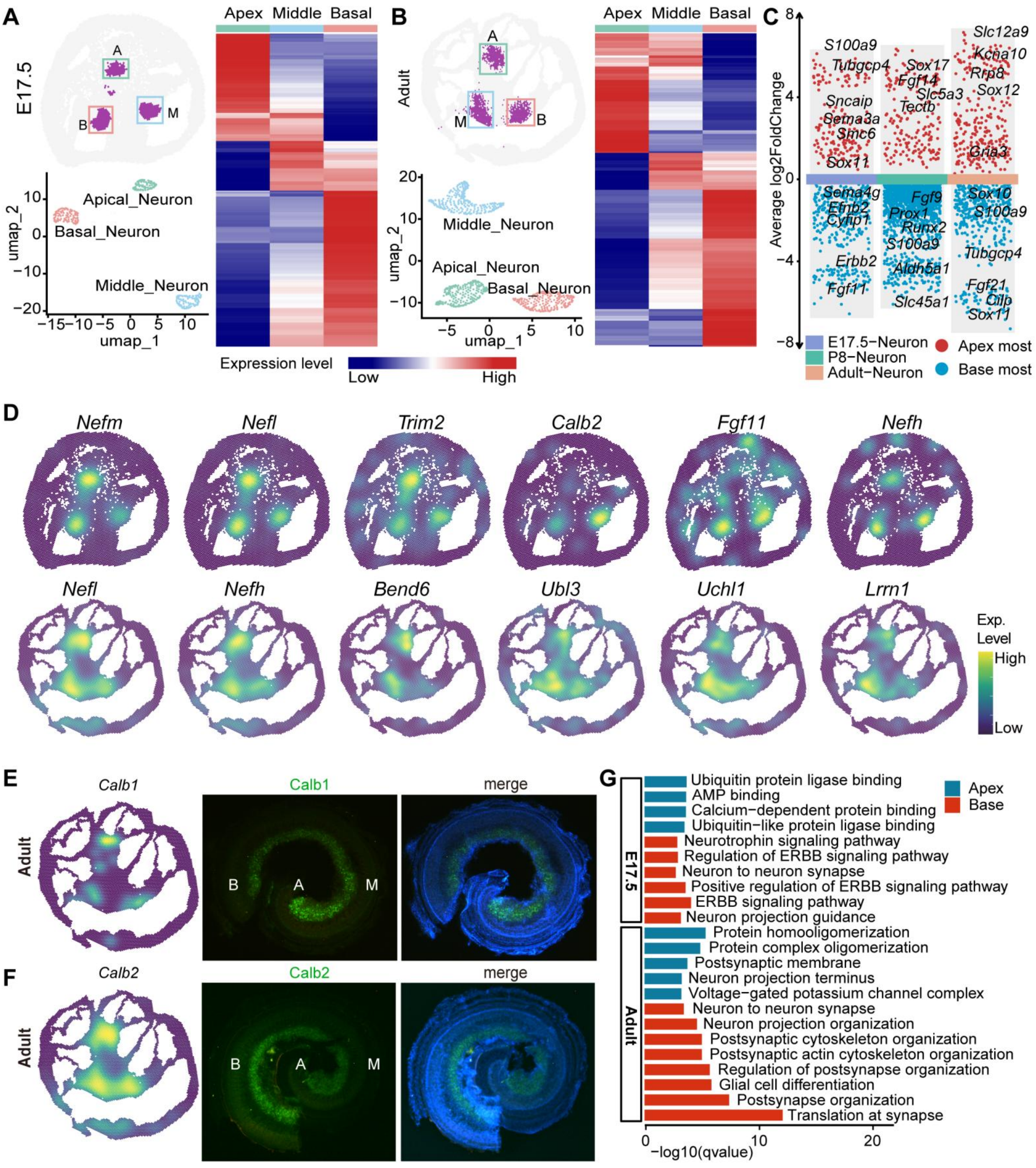

A
E17.5
A
B
M
Apical_Neuron
Basal_Neuron
Middle_Neuron
umap_1
umap_2
Apex
Middle
Basal
B
Adult
A
M
B
Middle_Neuron
Apical_Neuron
Basal_Neuron
Expression level
Low
High
C
Average log2FoldChange
S100a9
Tubgcp4
Sncaip
Sema3a
Smc6
Sox11
Sema4g
Efnb2
Cyfip1
Erbb2
Fgf11
Sox17
Fgf14
Slc5a3
Tectb
Fgf9
Prox1
Runx2
S100a9
Aldh5a1
Slc45a1
Slc12a9
Kcna10
Rrp8
Sox12
Gria3
Sox10
S100a9
Tubgcp4
Fgf21
Cilp
Sox11
E17.5−Neuron
P8−Neuron
Adult−Neuron
Apex most
Base most
D
Nefm
Nefl
Trim2
Calb2
Fgf11
Nefh
Nefl
Nefh
Bend6
Ubl3
Uchl1
Lrrn1
Exp. Level
High
Low
E
Calb1
Adult
Calb1
merge
B
A
M
F
Calb2
Adult
Calb2
merge
B
A
M
G
Apex
Base
E17.5
Ubiquitin protein ligase binding
AMP binding
Calcium−dependent protein binding
Ubiquitin−like protein ligase binding
Neurotrophin signaling pathway
Regulation of ERBB signaling pathway
Neuron to neuron synapse
Positive regulation of ERBB signaling pathway
ERBB signaling pathway
Neuron projection guidance
Adult
Protein homooligomerization
Protein complex oligomerization
Postsynaptic membrane
Neuron projection terminus
Voltage−gated potassium channel complex
Neuron to neuron synapse
Neuron projection organization
Postsynaptic cytoskeleton organization
Postsynaptic actin cytoskeleton organization
Regulation of postsynapse organization
Glial cell differentiation
Postsynapse organization
Translation at synapse
−log10(qvalue)

**S3-2 Fig**

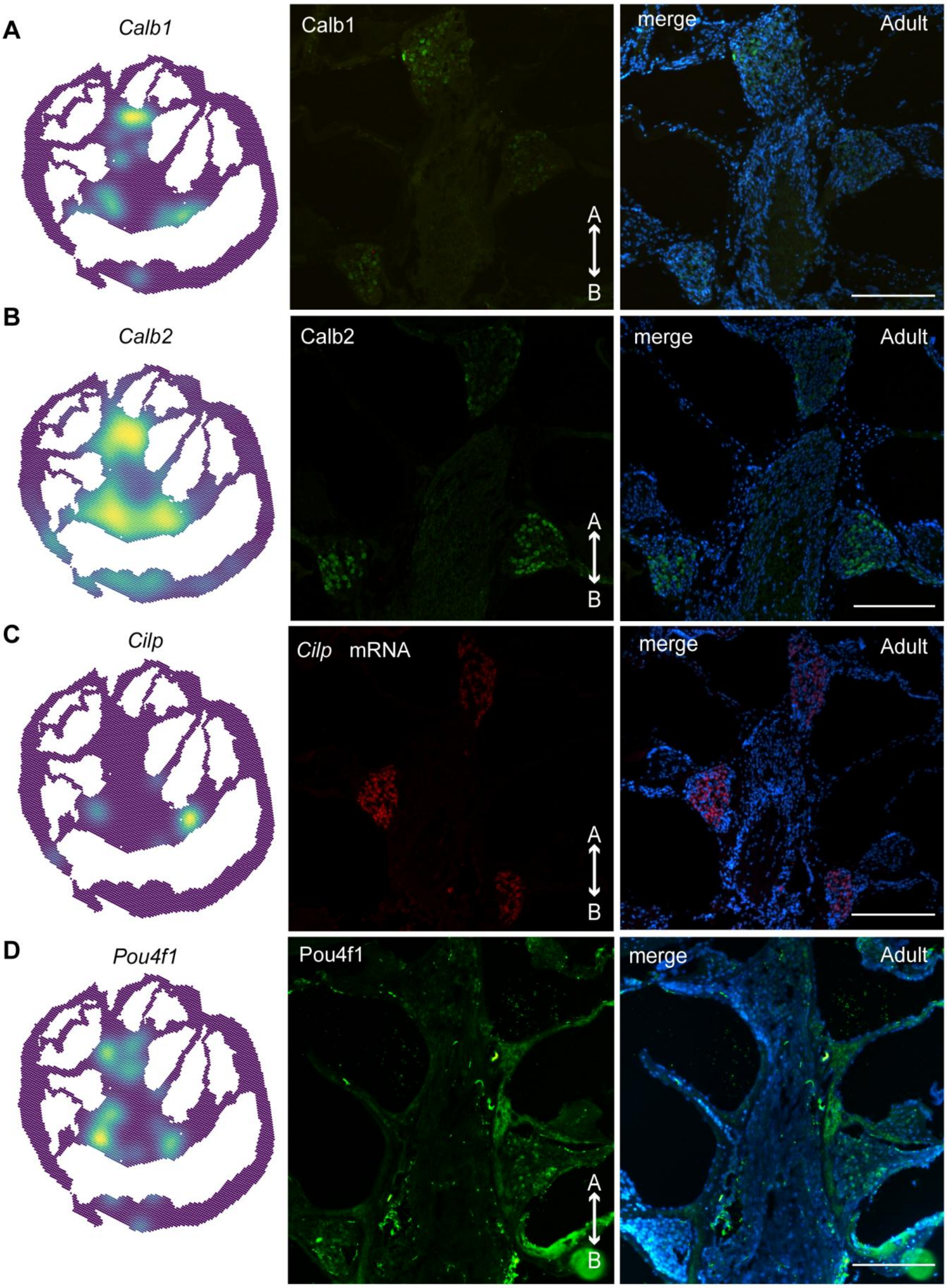
A
Calb1
Calb1
merge
Adult
A
B
B
Calb2
Calb2
merge
Adult
A
B
C
Cilp
Cilp mRNA
merge
Adult
A
B
D
Pou4f1
Pou4f1
merge
Adult
A
B

S4 Fig

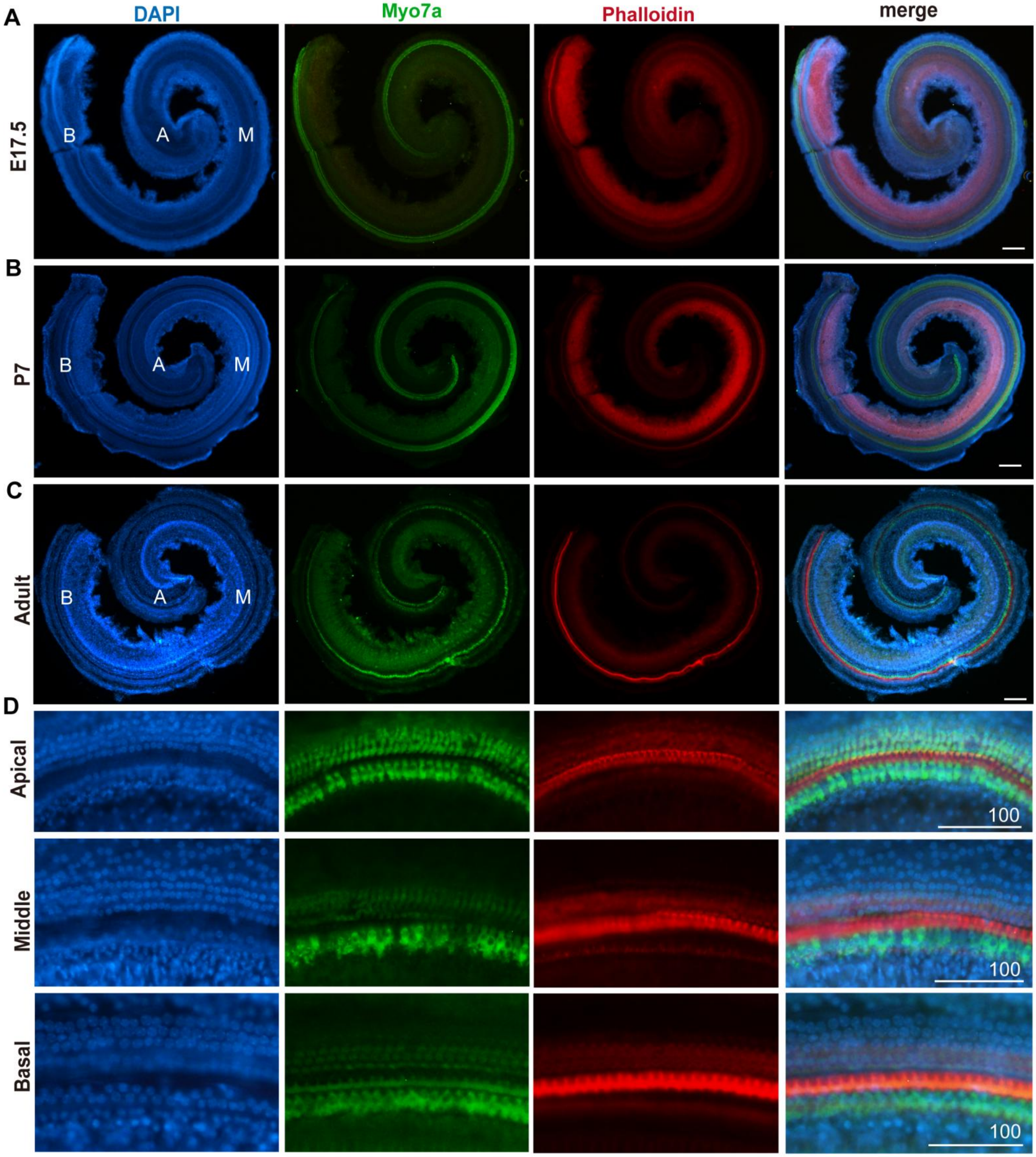

A
B
C
D
DAPI
Myo7a
Phalloidin
merge
E17.5
P7
Adult
Apical
Middle
Basal
B
A
M
100

**S5-1 Fig**

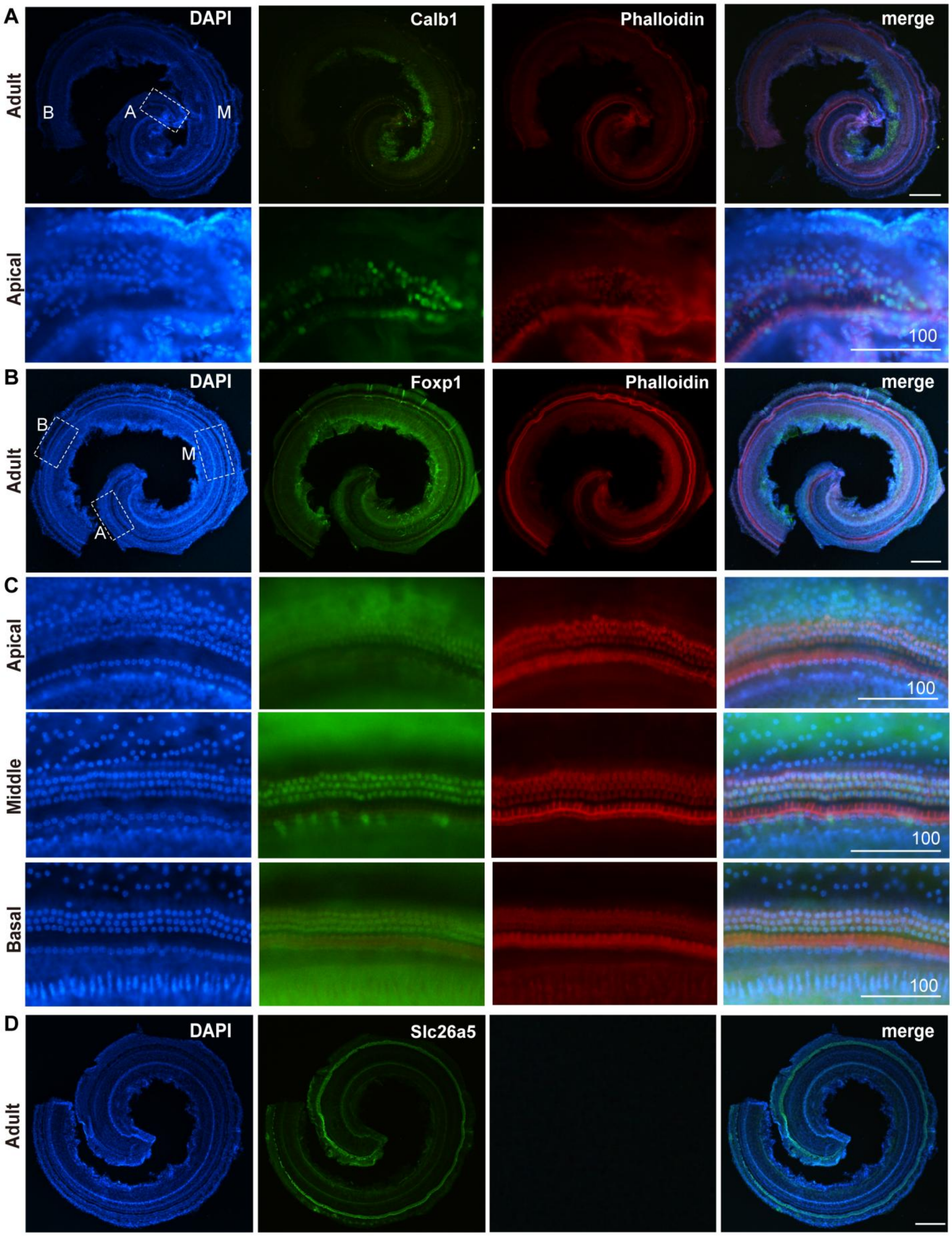

A
Adult
DAPI
B
A
M
Calb1
Phalloidin
merge
Apical
100
B
Adult
DAPI
B
M
A
Foxp1
Phalloidin
merge
C
Apical
100
Middle
100
Basal
100
D
Adult
DAPI
Slc26a5
merge

**S5-2 Fig**

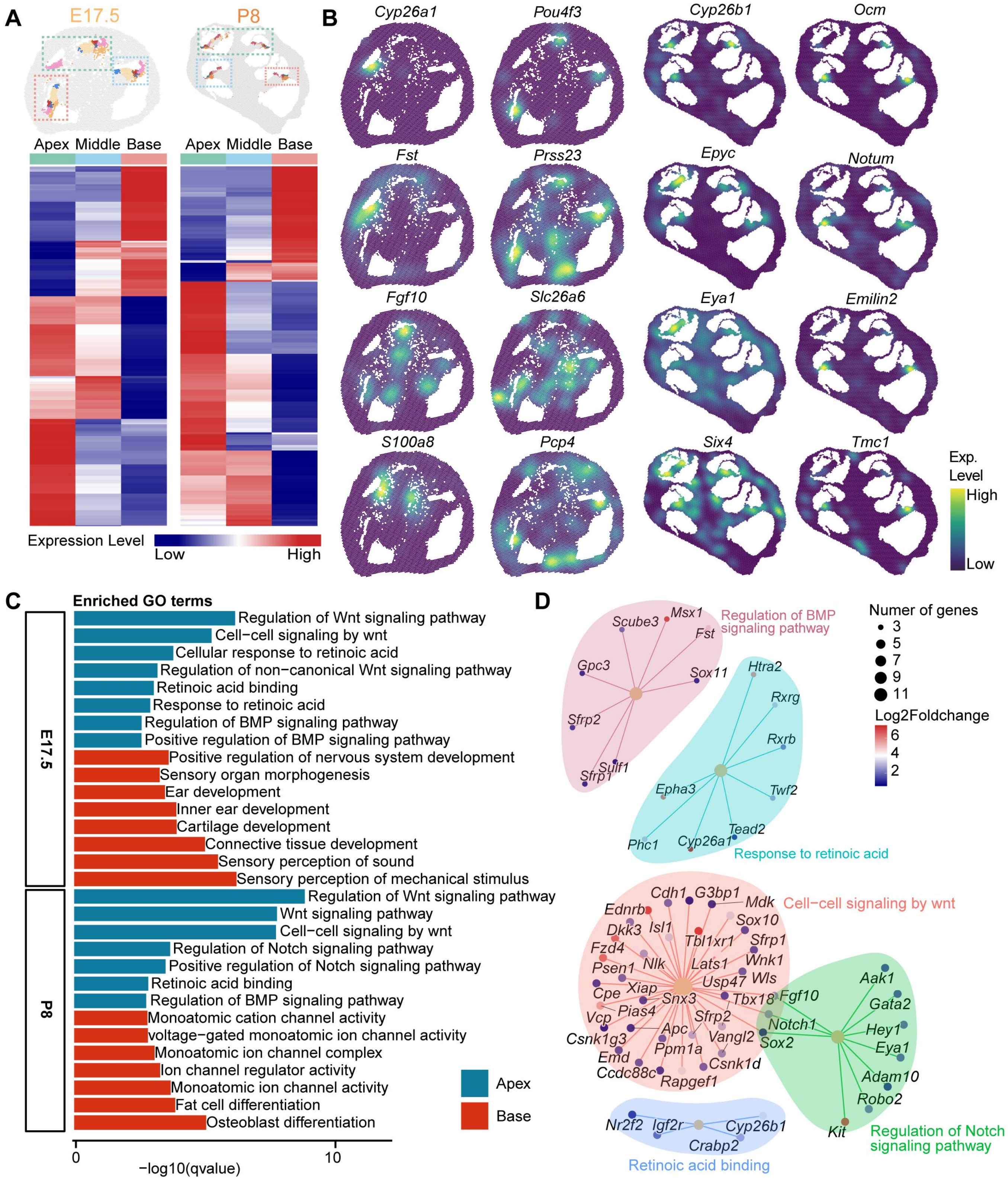
A
E17.5
P8
Apex Middle Base
Apex Middle Base
Expression Level
Low
High
B
Cyp26a1
Pou4f3
Cyp26b1
Ocm
Fst
Prss23
Epyc
Notum
Fgf10
Slc26a6
Eya1
Emilin2
S100a8
Pcp4
Six4
Tmc1
Exp. Level
High
Low
C
Enriched GO terms
E17.5
Regulation of Wnt signaling pathway
Cell−cell signaling by wnt
Cellular response to retinoic acid
Regulation of non−canonical Wnt signaling pathway
Retinoic acid binding
Response to retinoic acid
Regulation of BMP signaling pathway
Positive regulation of BMP signaling pathway
Positive regulation of nervous system development
Sensory organ morphogenesis
Ear development
Inner ear development
Cartilage development
Connective tissue development
Sensory perception of sound
Sensory perception of mechanical stimulus
P8
Regulation of Wnt signaling pathway
Wnt signaling pathway
Cell−cell signaling by wnt
Regulation of Notch signaling pathway
Positive regulation of Notch signaling pathway
Retinoic acid binding
Regulation of BMP signaling pathway
Monoatomic cation channel activity
voltage−gated monoatomic ion channel activity
Monoatomic ion channel complex
Ion channel regulator activity
Monoatomic ion channel activity
Fat cell differentiation
Osteoblast differentiation
Apex
Base
0
10
−log10(qvalue)
D
Regulation of BMP signaling pathway
Scube3 Msx1 Fst Gpc3 Sox11 Sfrp2 Sulf1 Sfrp1
Response to retinoic acid
Htra2 Rxrg Rxrb Epha3 Twf2 Tead2 Phc1 Cyp26a1
Cell−cell signaling by wnt
Cdh1 G3bp1 Mdk Ednrb Sox10 Dkk3 Isl1 Tbl1xr1 Sfrp1 Fzd4 Wnk1 Psen1 Nlk Lats1 Usp47 Wls Cpe Xiap Snx3 Tbx18 Vcp Pias4 Sfrp2 Csnk1g3 Apc Vangl2 Sox2 Emd Ppm1a Csnk1d Ccdc88c Rapgef1
Regulation of Notch signaling pathway
Fgf10 Aak1 Gata2 Notch1 Hey1 Eya1 Adam10 Robo2 Kit
Retinoic acid binding
Nr2f2 Igf2r Cyp26b1 Crabp2
Numer of genes
3
5
7
9
11
Log2Foldchange
6
4
2

#### S5-3 Fig

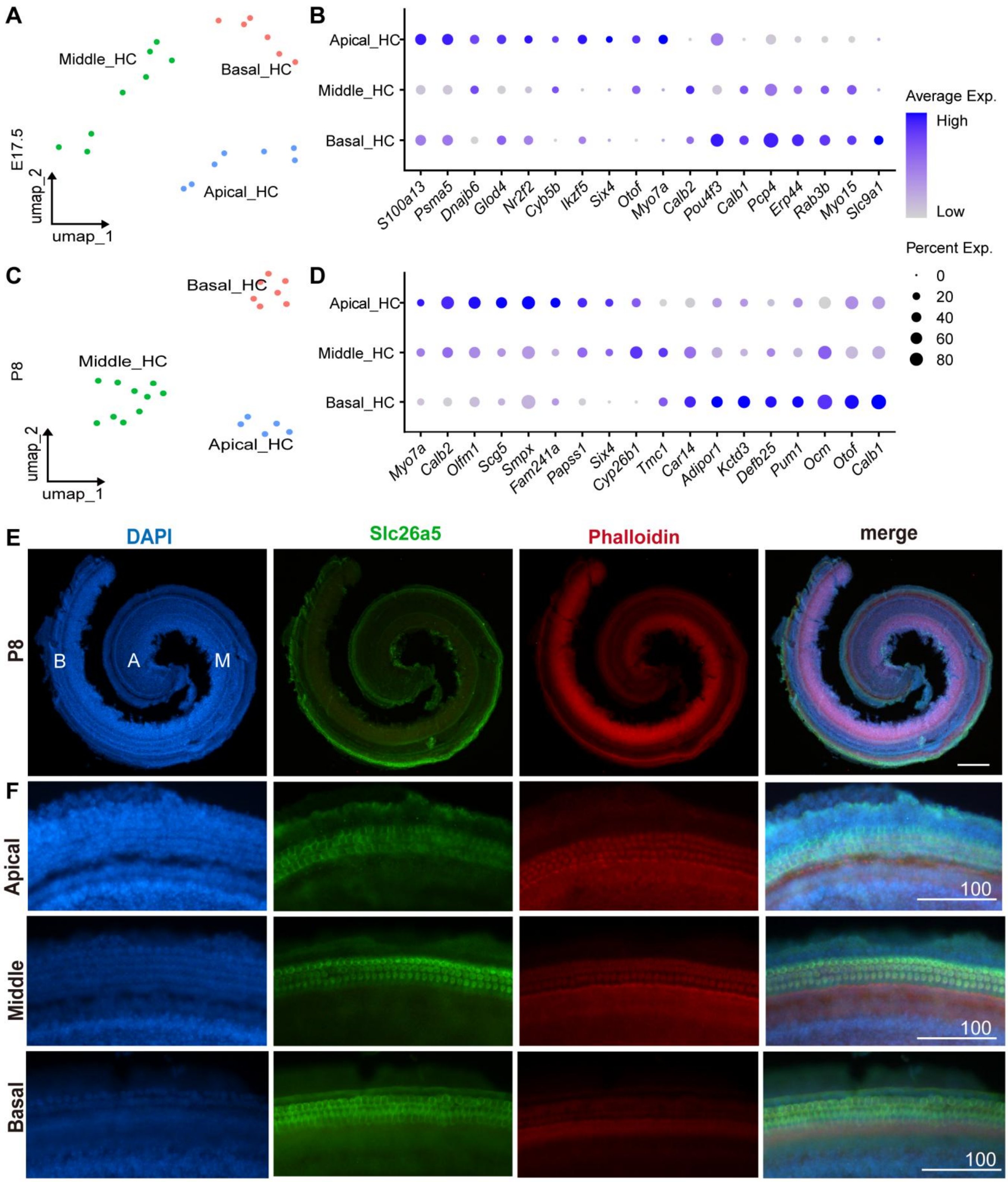

A
B
C
D
E
F
Middle_HC
Basal_HC
Apical_HC
E17.5
umap_2
umap_1
P8
Average Exp.
High
Low
Percent Exp.
0
20
40
60
80
S100a13
Psma5
Dnajb6
Glod4
Nr2f2
Cyb5b
Ikzf5
Six4
Otof
Myo7a
Calb2
Pou4f3
Calb1
Pcp4
Erp44
Rab3b
Myo15
Slc9a1
Olfm1
Scg5
Smpx
Fam241a
Papss1
Cyp26b1
Tmc1
Car14
Adipor1
Kctd3
Defb25
Pum1
Ocm
DAPI
Slc26a5
Phalloidin
merge
B
A
M
Apical
Middle
Basal
100

#### S6-1 Fig

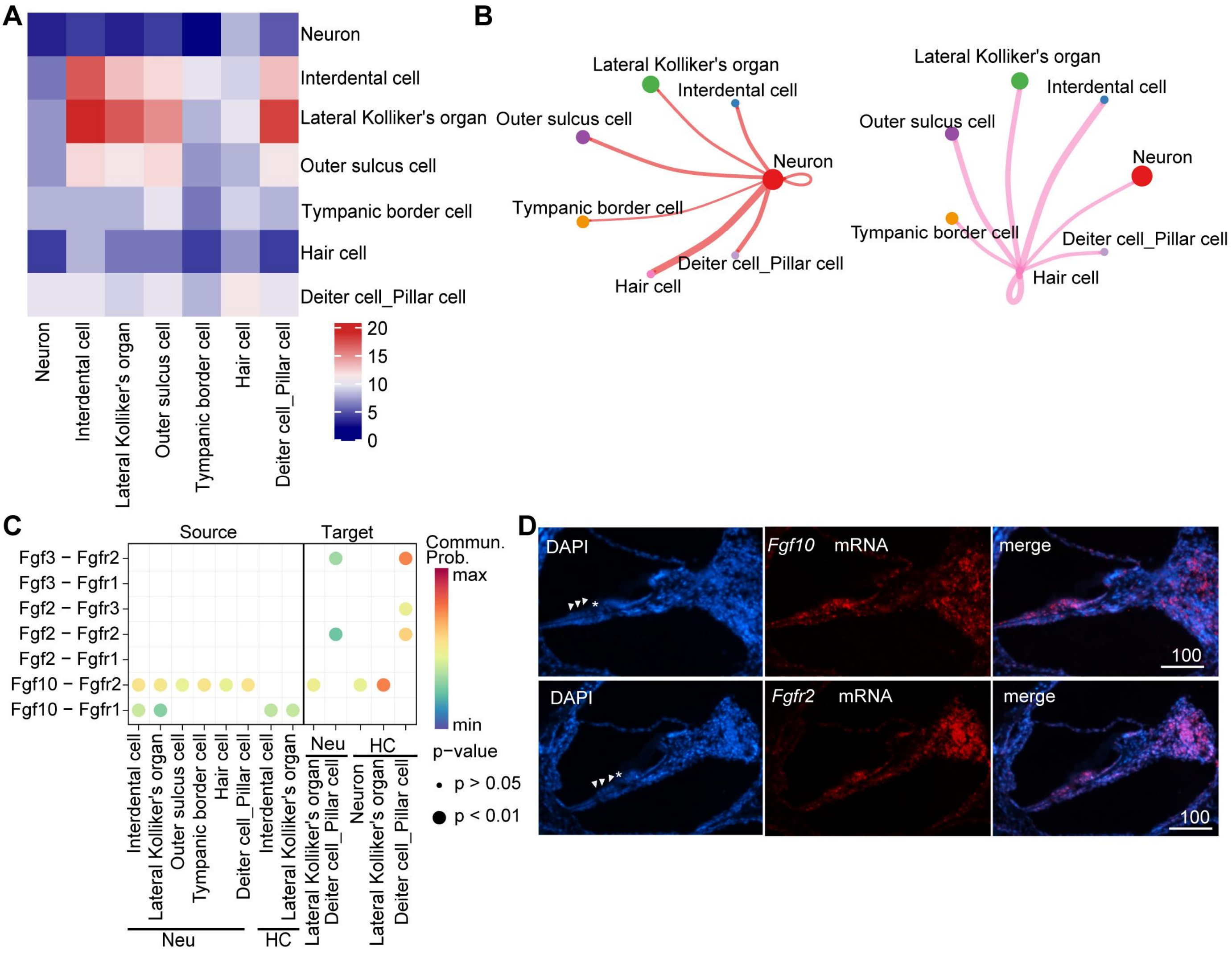
A
Neuron
Interdental cell
Lateral Kolliker's organ
Outer sulcus cell
Tympanic border cell
Hair cell
Deiter cell_Pillar cell
20
15
10
5
0
B
Lateral Kolliker's organ
Interdental cell
Outer sulcus cell
Neuron
Tympanic border cell
Deiter cell_Pillar cell
Hair cell
C
Source
Target
Fgf3 − Fgfr2
Fgf3 − Fgfr1
Fgf2 − Fgfr3
Fgf2 − Fgfr2
Fgf2 − Fgfr1
Fgf10 − Fgfr2
Fgf10 − Fgfr1
Commun. Prob.
max
min
p−value
p > 0.05
p < 0.01
Neu
HC
D
DAPI
Fgf10 mRNA
merge
100
DAPI
Fgfr2 mRNA
merge
100

### S6-2 Fig

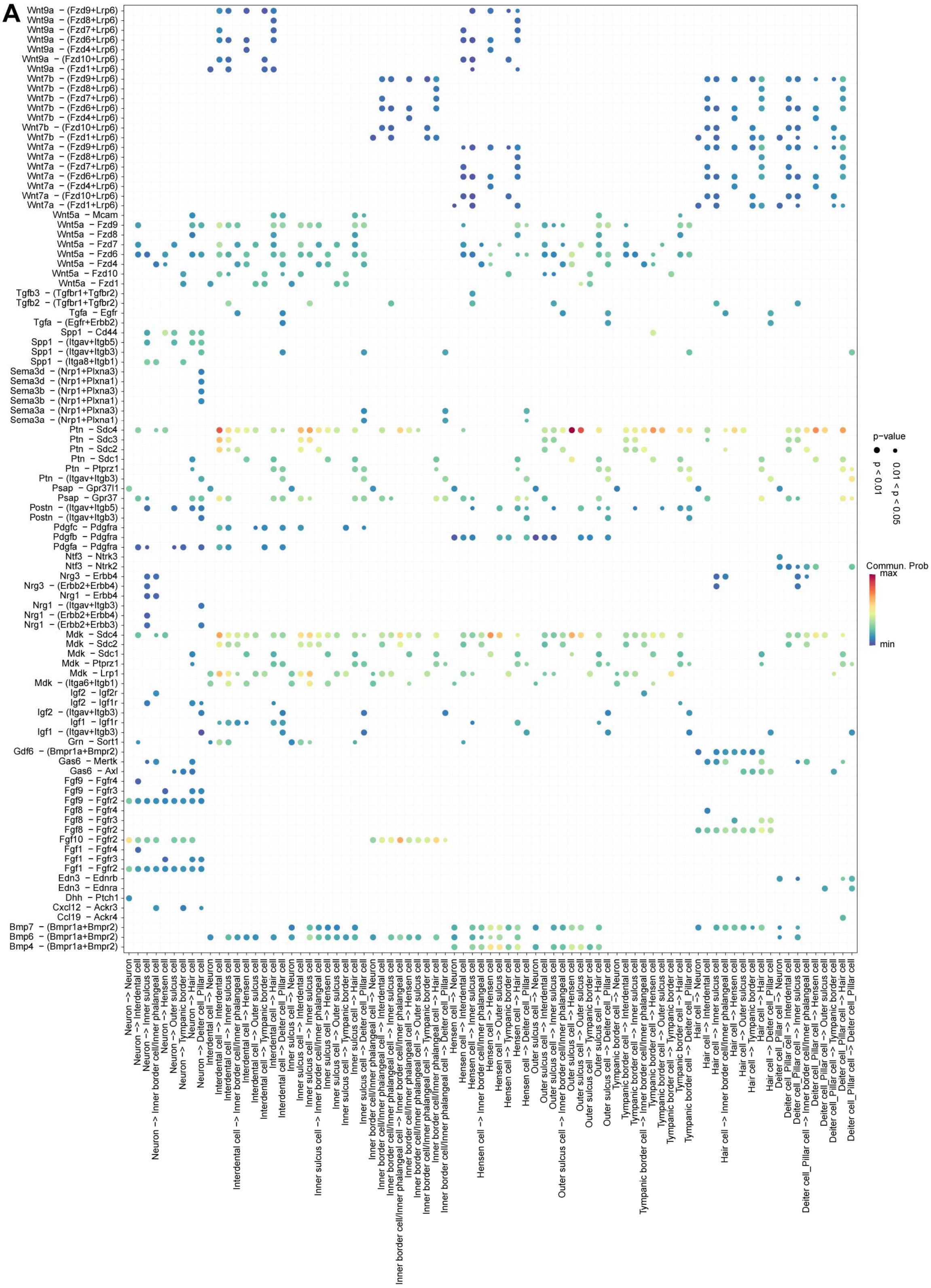

#### S7-1 Fig

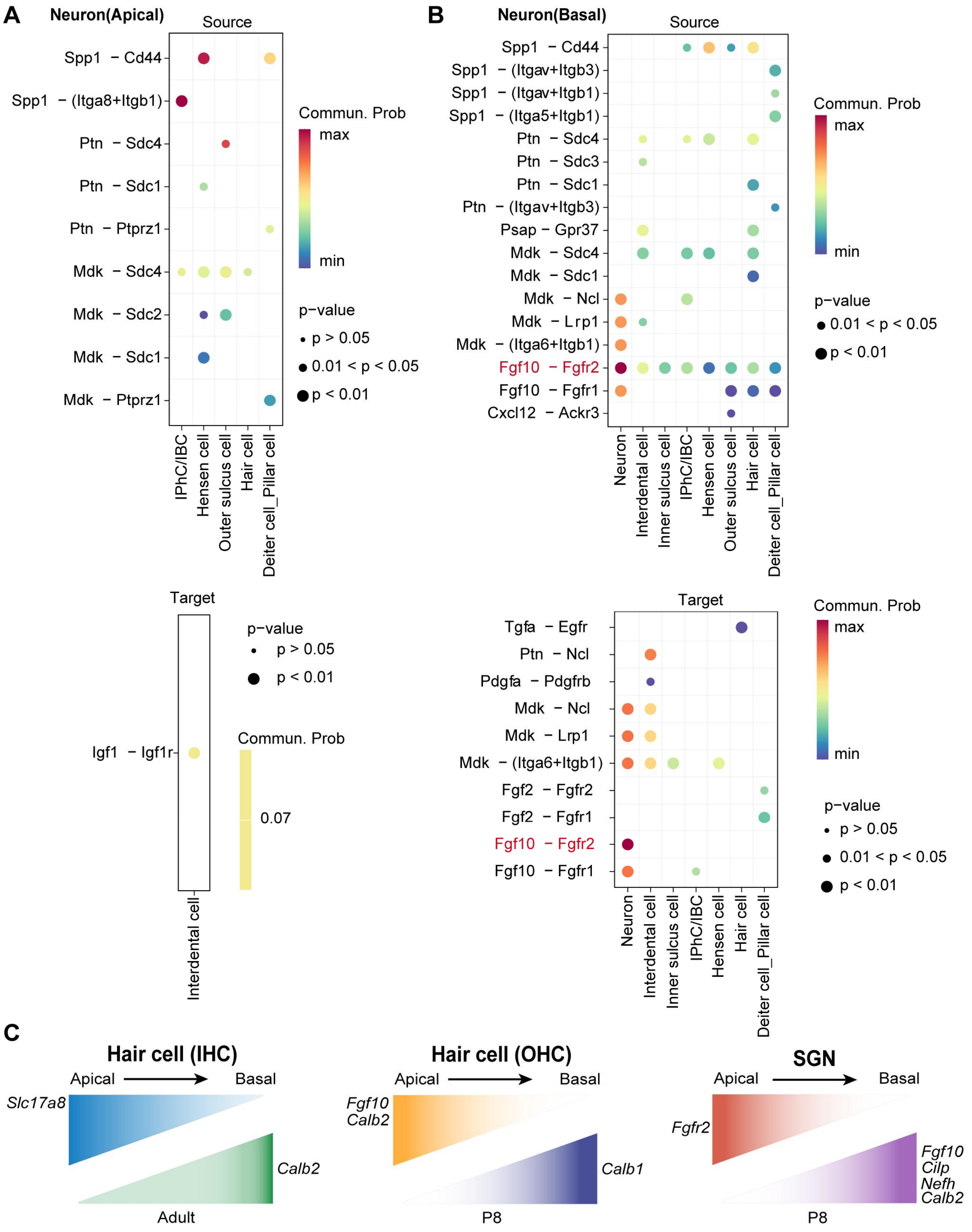
A
Neuron(Apical)
Source
Spp1 − Cd44
Spp1 − (Itga8+Itgb1)
Ptn − Sdc4
Ptn − Sdc1
Ptn − Ptprz1
Mdk − Sdc4
Mdk − Sdc2
Mdk − Sdc1
Mdk − Ptprz1
IPhC/IBC
Hensen cell
Outer sulcus cell
Hair cell
Deiter cell_Pillar cell
Commun. Prob
max
min
p−value
p > 0.05
0.01 < p < 0.05
p < 0.01
B
Neuron(Basal)
Source
Spp1 − Cd44
Spp1 − (Itgav+Itgb3)
Spp1 − (Itgav+Itgb1)
Spp1 − (Itga5+Itgb1)
Ptn − Sdc4
Ptn − Sdc3
Ptn − Sdc1
Ptn − (Itgav+Itgb3)
Psap − Gpr37
Mdk − Sdc4
Mdk − Sdc1
Mdk − Ncl
Mdk − Lrp1
Mdk − (Itga6+Itgb1)
Fgf10 − Fgfr2
Fgf10 − Fgfr1
Cxcl12 − Ackr3
Neuron
Interdental cell
Inner sulcus cell
IPhC/IBC
Hensen cell
Outer sulcus cell
Hair cell
Deiter cell_Pillar cell
Commun. Prob
max
min
p−value
0.01 < p < 0.05
p < 0.01
Target
Igf1 − Igf1r
Interdental cell
p−value
p > 0.05
p < 0.01
Commun. Prob
0.07
Target
Tgfa − Egfr
Ptn − Ncl
Pdgfa − Pdgfrb
Mdk − Ncl
Mdk − Lrp1
Mdk − (Itga6+Itgb1)
Fgf2 − Fgfr2
Fgf2 − Fgfr1
Fgf10 − Fgfr2
Fgf10 − Fgfr1
Neuron
Interdental cell
Inner sulcus cell
IPhC/IBC
Hensen cell
Hair cell
Deiter cell_Pillar cell
Commun. Prob
max
min
p−value
p > 0.05
0.01 < p < 0.05
p < 0.01
C
Hair cell (IHC)
Apical
Basal
Slc17a8
Calb2
Adult
Hair cell (OHC)
Apical
Basal
Fgf10
Calb2
Calb1
P8
SGN
Apical
Basal
Fgfr2
Fgf10
Cilp
Nefh
Calb2
P8

#### S7-2 Fig

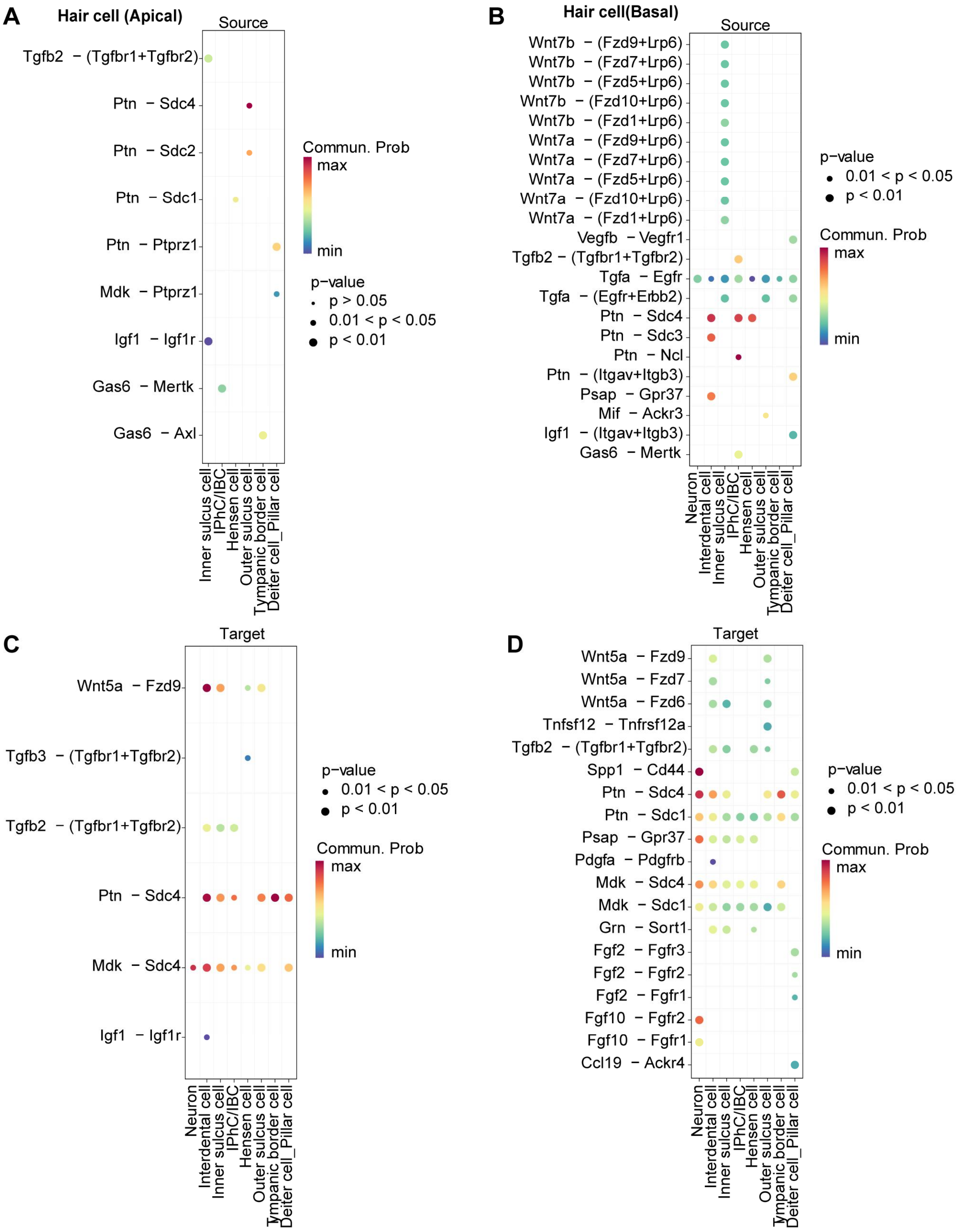

A
Hair cell (Apical)
Source
Tgfb2 − (Tgfbr1+Tgfbr2)
Ptn − Sdc4
Ptn − Sdc2
Ptn − Sdc1
Ptn − Ptprz1
Mdk − Ptprz1
Igf1 − Igf1r
Gas6 − Mertk
Gas6 − Axl
Inner sulcus cell
IPhC/IBC
Hensen cell
Outer sulcus cell
Tympanic border cell
Deiter cell_Pillar cell
Commun. Prob
max
min
p−value
p > 0.05
0.01 < p < 0.05
p < 0.01
B
Hair cell(Basal)
Source
Wnt7b − (Fzd9+Lrp6)
Wnt7b − (Fzd7+Lrp6)
Wnt7b − (Fzd5+Lrp6)
Wnt7b − (Fzd10+Lrp6)
Wnt7b − (Fzd1+Lrp6)
Wnt7a − (Fzd9+Lrp6)
Wnt7a − (Fzd7+Lrp6)
Wnt7a − (Fzd5+Lrp6)
Wnt7a − (Fzd10+Lrp6)
Wnt7a − (Fzd1+Lrp6)
Vegfb − Vegfr1
Tgfb2 − (Tgfbr1+Tgfbr2)
Tgfa − Egfr
Tgfa − (Egfr+Erbb2)
Ptn − Sdc4
Ptn − Sdc3
Ptn − Ncl
Ptn − (Itgav+Itgb3)
Psap − Gpr37
Mif − Ackr3
Igf1 − (Itgav+Itgb3)
Gas6 − Mertk
Neuron
Interdental cell
Inner sulcus cell
IPhC/IBC
Hensen cell
Outer sulcus cell
Tympanic border cell
Deiter cell_Pillar cell
p−value
0.01 < p < 0.05
p < 0.01
Commun. Prob
max
min
C
Target
Wnt5a − Fzd9
Tgfb3 − (Tgfbr1+Tgfbr2)
Tgfb2 − (Tgfbr1+Tgfbr2)
Ptn − Sdc4
Mdk − Sdc4
Igf1 − Igf1r
Neuron
Interdental cell
Inner sulcus cell
IPhC/IBC
Hensen cell
Outer sulcus cell
Tympanic border cell
Deiter cell_Pillar cell
p−value
0.01 < p < 0.05
p < 0.01
Commun. Prob
max
min
D
Target
Wnt5a − Fzd9
Wnt5a − Fzd7
Wnt5a − Fzd6
Tnfsf12 − Tnfrsf12a
Tgfb2 − (Tgfbr1+Tgfbr2)
Spp1 − Cd44
Ptn − Sdc4
Ptn − Sdc1
Psap − Gpr37
Pdgfa − Pdgfrb
Mdk − Sdc4
Mdk − Sdc1
Grn − Sort1
Fgf2 − Fgfr3
Fgf2 − Fgfr2
Fgf2 − Fgfr1
Fgf10 − Fgfr2
Fgf10 − Fgfr1
Ccl19 − Ackr4
Neuron
Interdental cell
Inner sulcus cell
IPhC/IBC
Hensen cell
Outer sulcus cell
Tympanic border cell
Deiter cell_Pillar cell
p−value
0.01 < p < 0.05
p < 0.01
Commun. Prob
max
min

**S7-3 Fig**

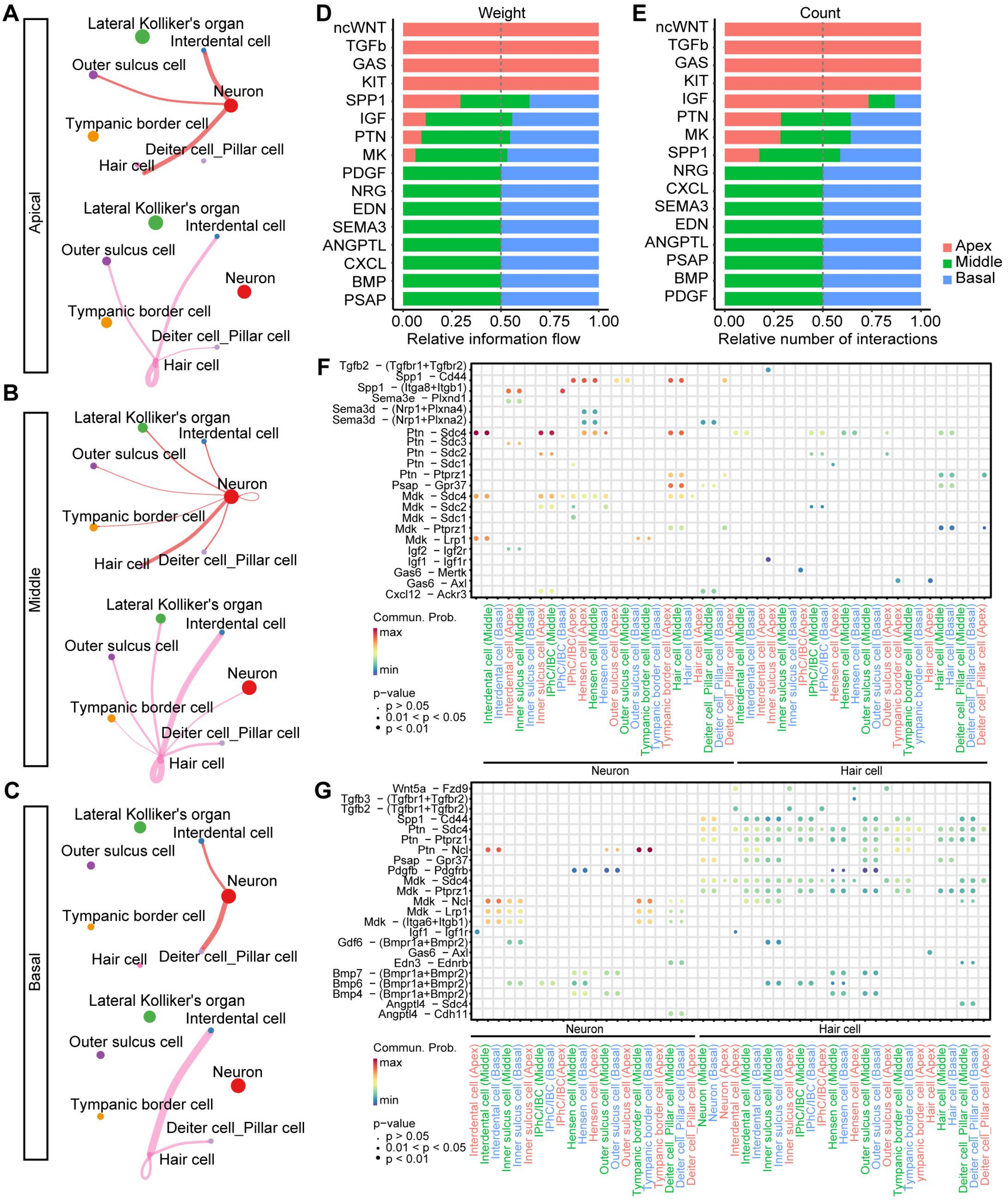

A
Apical
Lateral Kolliker's organ
Interdental cell
Outer sulcus cell
Neuron
Tympanic border cell
Deiter cell_Pillar cell
Hair cell
B
Middle
C
Basal
D
Weight
Relative information flow
E
Count
Relative number of interactions
Apex
Middle
Basal
F
G
Commun. Prob.
max
min
p-value
p > 0.05
0.01 < p < 0.05
p < 0.01
Neuron
Hair cell

**S8 Fig**

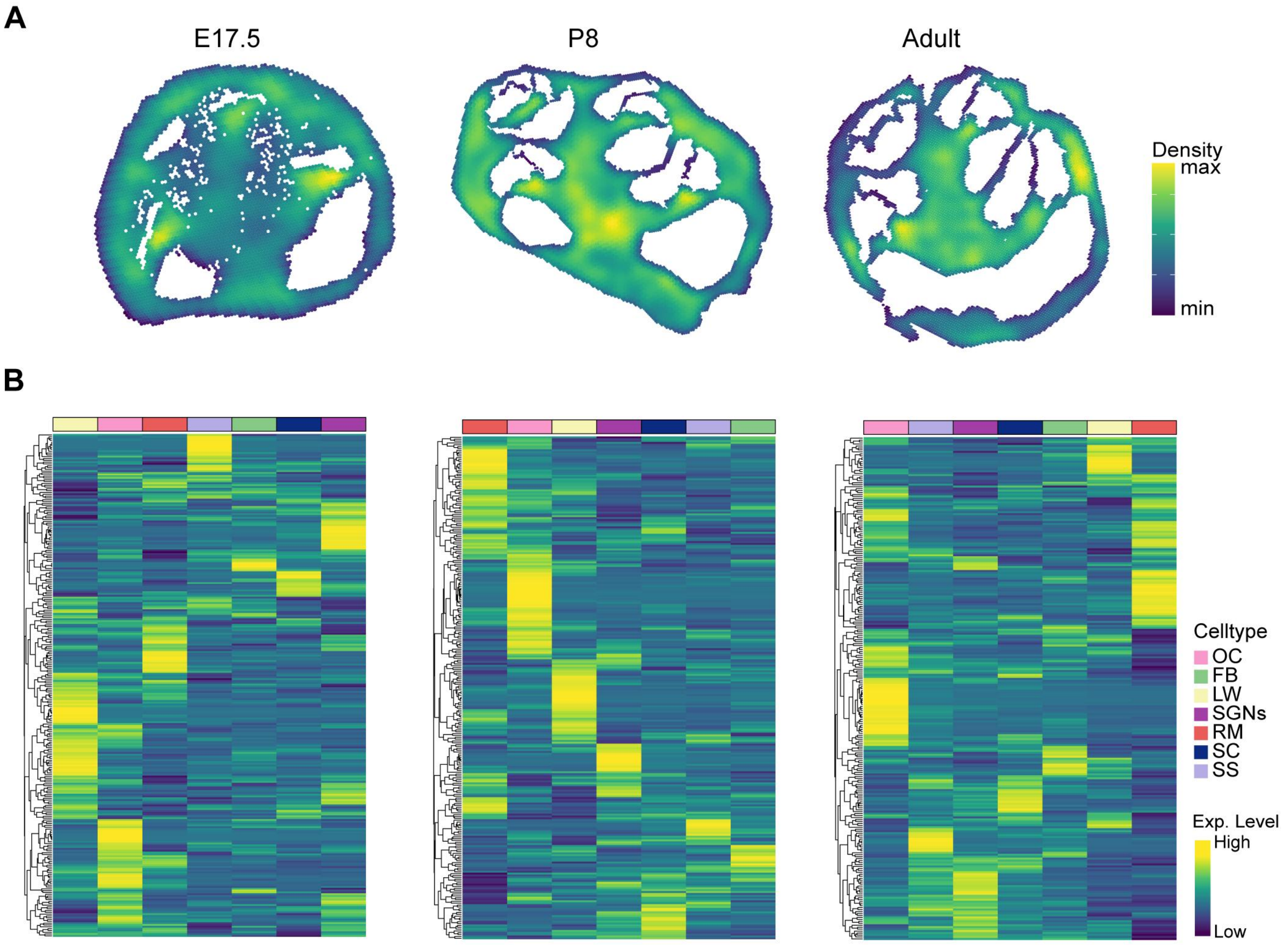
A
E17.5
P8
Adult
Density
max
min
B
Celltype
OC
FB
LW
SGNs
RM
SC
SS
Exp. Level
High
Low
